# Spatial and temporal localization of immune transcripts defines hallmarks and diversity in the tuberculosis granuloma

**DOI:** 10.1101/509547

**Authors:** Berit Carow, Thomas Hauling, Xiaoyan Qian, Igor Kramnik, Mats Nilsson, Martin E Rottenberg

## Abstract

Granulomas are the pathological hallmark of Tuberculosis (TB), and the niche in which bacilli can either grow and disseminate or the immunological microenvironment in which host cells interact to prevent bacterial dissemination. Here, after in situ sequencing, thirty-four immune transcripts in lung sections from *Mycobacterium tuberculosis*-infected mice were aligned to the tissue morphology at cellular resolution, allowing the analysis of local immune interactions in the granuloma.

Co-localizing transcript networks at <10 μm in C57BL/6 mouse granulomas increased in complexity with time after infection. B-cell clusters developed late after infection. Transcripts from activated macrophages were enriched at subcellular distances from *M. tuberculosis*. Encapsulated C3HeB/FeJ granulomas showed necrotic centers with transcripts associated with immunosuppression (*foxp3, il10*), while those in the granuloma rims associated with activated T cells and macrophages. Highly diverse networks with common interactors were observed in similar lesions.

Thus, different immune landscapes of *M. tuberculosis* granulomas depending on the time after infection, the histopathological features of the lesion and the proximity to bacteria were here defined.

## Main

Tuberculosis (TB), caused by infection with *Mycobacterium tuberculosis,* remains a leading public health problem worldwide. In 2016, 10.4 million people developed TB and 1.7 million patients died from the disease^1^. The challenges to control TB are enormous: resistance to available drugs against *M. tuberculosis* is increasing, while the only available vaccine against TB is only partially protective. One quarter of the global population is latently infected with *M. tuberculosis* with a 10% probability of those infected to develop active TB during their lifetime. To understand why reactivation of *M. tuberculosis* occurs in some but not in other individuals is central for development of new vaccines and therapies.

Infection occurs when the inhaled *M. tuberculosis* are phagocytized by resident lung alveolar macrophages^2^. Infected cells recruit mononuclear phagocytes to the infection site, forming a nascent granuloma. During the subclinical stage of infection, the granuloma provides the immune environment required for the containment of bacteria. *M. tuberculosis*-specific T cells are crucial for the granuloma maturation, maintenance and control of the bacterial spread^3^. However, if due to impaired immunity the integrity of the granuloma is lost, reactivation of *M. tuberculosis* leads to the destruction of the lung structure and to the transmission of *M. tuberculosis* to other humans.

In association with the diverse outcome of infection, studies from autopsies have shown important diversity in the granuloma histology. In addition to the encapsulated granuloma with a caseous necrotic center, TB granulomas can be non-necrotizing, neutrophil-rich, mineralized, fibrotic or cavitary^4,5^. TB granulomas in commonly used inbred mouse strains such as C57BL/6 do not develop necrotizing lesions, while encapsulated necrotizing granulomas are found in other strains such as the C3HeB/FeJ^6–8^. While the histological features of granulomas have been well characterized, the immune and inflammatory mechanisms that underlie variable granuloma dynamics and clinical outcomes of TB infection remain to be further elucidated.

In order to understand the immunological architecture of the murine TB granulomas, we have used a novel method for highly sensitive multiplexed *in situ* imaging of selected immune mRNA species. The method is based on rolling-circle amplification (RCA) of padlock probes and on sequencing-by-ligation chemistry, as reported^9^. RCA has been used to produce highly specific amplified products that enabled detection of individual mRNA molecules *in situ* in the unperturbed context of fixed tissues at cellular resolution^9,10^.

We show that granulomas developed with time showing increasing complexity and diversity of co-expressing molecular networks. Transcripts corresponding to activated macrophages localized at subcellular distances from *M. tuberculosis*. Moreover, we describe profound differences between different granuloma regions and between encapsulated necrotic and non-encapsulated granulomas. Common networks underlying the observed heterogeneity could be defined for different areas of the granulomas, for granulomas at different time points and from different mouse strains.

## Results

### Specificity, reproducibility and performance of in situ sequencing

The *in situ* sequencing technique was used to localize simultaneously 34 immune transcripts coding for chemokine receptors, cytokines, effector molecules and surface molecules that define immune populations in paraformaldehyde-fixed sections of lungs from *M. tuberculosis*-infected mice. Three consecutive sections from C57BL/6 mice at 3, 8 and 12 weeks after aerosol infection with *M. tuberculosis* (wpi) and those from C3HeB/FeJ mice that develop encapsulated necrotic granulomas were used^11^. Transcripts were aligned with the histopathological features of the same lung section (Figure S1).

Non-specific signals (when base calling did not correspond to those built-in the barcoding sequences) were minimized by increasing the signal threshold, while the density of specific signals remained mostly unaltered, indicating a high specificity of the reaction (Figure S2A). At the fixed threshold selected (0,45), the performance (the total number of signals) ranged from 7 to 15 x 10^4^ signals per section (Figure S2B). All transcripts were simultaneously and differentially detected in the lungs (Figure S2C). As expected, *cc10* mRNA, expressed by Clara cells, located in the epithelium of pulmonary airways, and *inos*, expressed by inflammatory cells, localized in the granuloma. These showed similar distribution in consecutive slides (Figure S2D). The ratio of the density of specific mRNAs in annotated lesions versus unaffected areas (defined in Figure S1) from consecutive sections were similar for some but the variance was higher for other transcripts (Figure S2E). This usually related to the higher sparsity of signals in the latter group.

### Maturation of the granuloma

Several immune transcripts located within the granulomas as shown by a signal and a density plot representation for the frequencies of *cd68, inos* and *cd3* mRNA (Figure 1A). All but 4 mRNA species (*cd4, cd40l, elane* and *il17a*) showed increased localization in the lesions compared to non-affected areas at 3 wpi, while all except *cd4* and *cd40l* preferentially located in the granuloma 12 wpi of C57BL/6 mice in consecutive slides (p<0.01 *χ*^2^ test).

**Figure 1.**
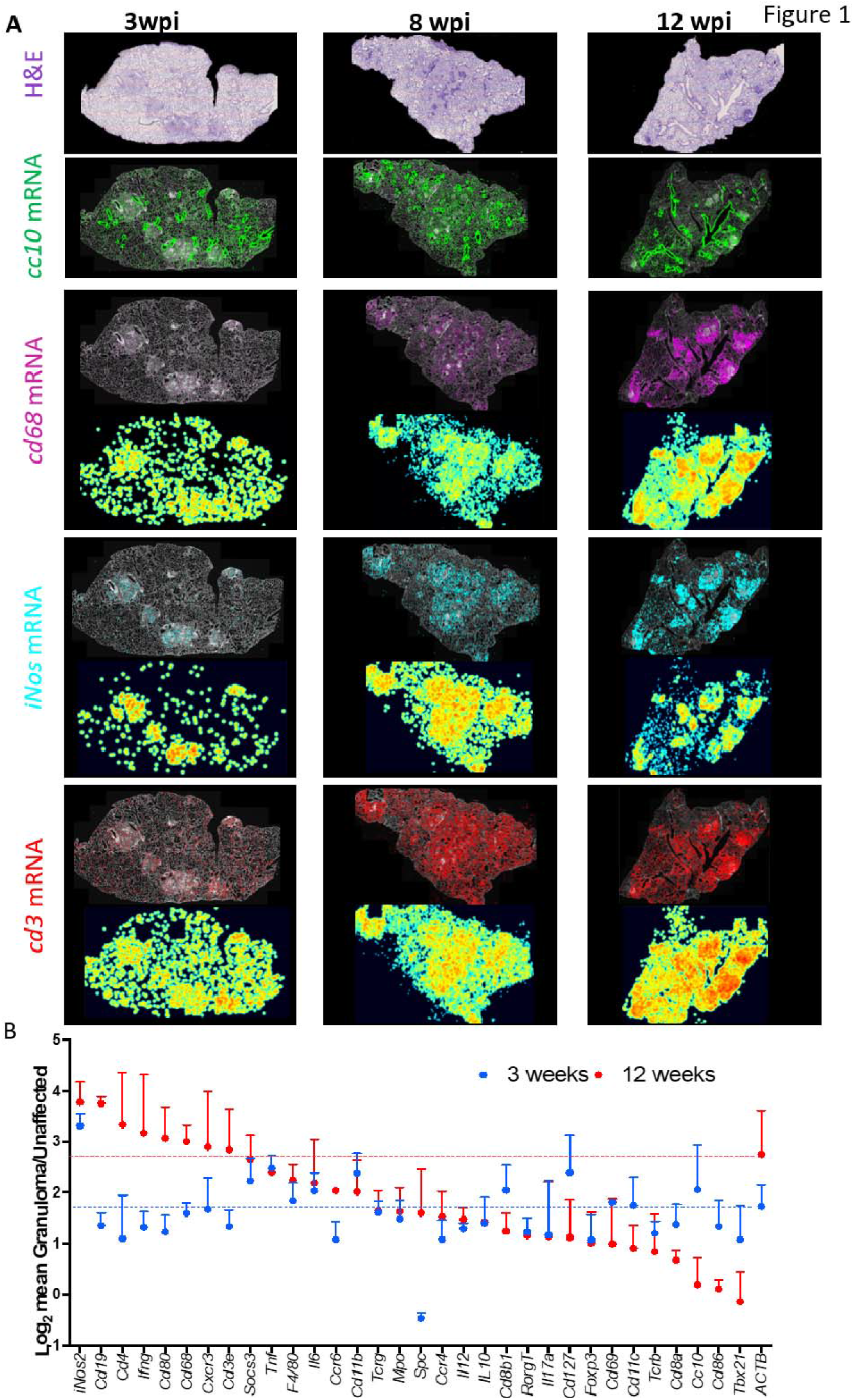
Preferential localization of immune sequences in granulomas during infection with M. _tuberculosis. A. At 3, 8 and 12 weeks post *M. tuberculosis* infection (wpi), the fixed lung tissue sections were stained with hematoxylin-eosin (H&E) and signals for *cc10, cd68, inos* and *cd3* were plotted on DAPI labelling as background. Pseudo-color density XY positional log_2_ plots of transcript representations are shown below *cd68, inos* and *cd3* transcripts. One representative of 3 consecutive sections is displayed. B. The ratio of amplified transcripts in granulomas vs unaffected regions was calculated for each transcript. The mean fold over unaffected region ± SEM in 3 consecutive sections is depicted. Sections from lungs at 3 and 12 weeks after infection are compared.

The house keeping *actb* mRNA was increased 2-fold in the granuloma compared to the unaffected regions, reflecting higher cellular density in the lesion (Figure S2E). Several transcripts in the lesions displayed higher relative frequencies than *actb* mRNA (Figure S2E). Several transcripts expressed by myeloid cells (i.e. *tnf, inos* and *il6*) localized at similar frequencies in granulomas from lungs at all time points after *M. tuberculosis* infection studied (Figure 1B). Instead, increased relative enrichment was observed for several transcripts associated with adaptive immune responses at 8 and 12 wpi (i.e. *cd19, ccr6, cd68, cxcr3* and *cd3e* mRNA) (Figure 1B).

We then analyzed transcripts co-expressed at <10 μm using the Insitunet app^12^. Transcripts with significant spatial co-expression were displayed as edges in a network map. Interactions were only studied when observed in at least 2 consecutive slides (Figure 2A). As an example, out of 561 possible mathematical combinations of pairs of the 34 transcripts at 8 wpi studied (C_34,2_), 19 transcript pairs significantly co-localized (Table S1).

**Figure 2.**
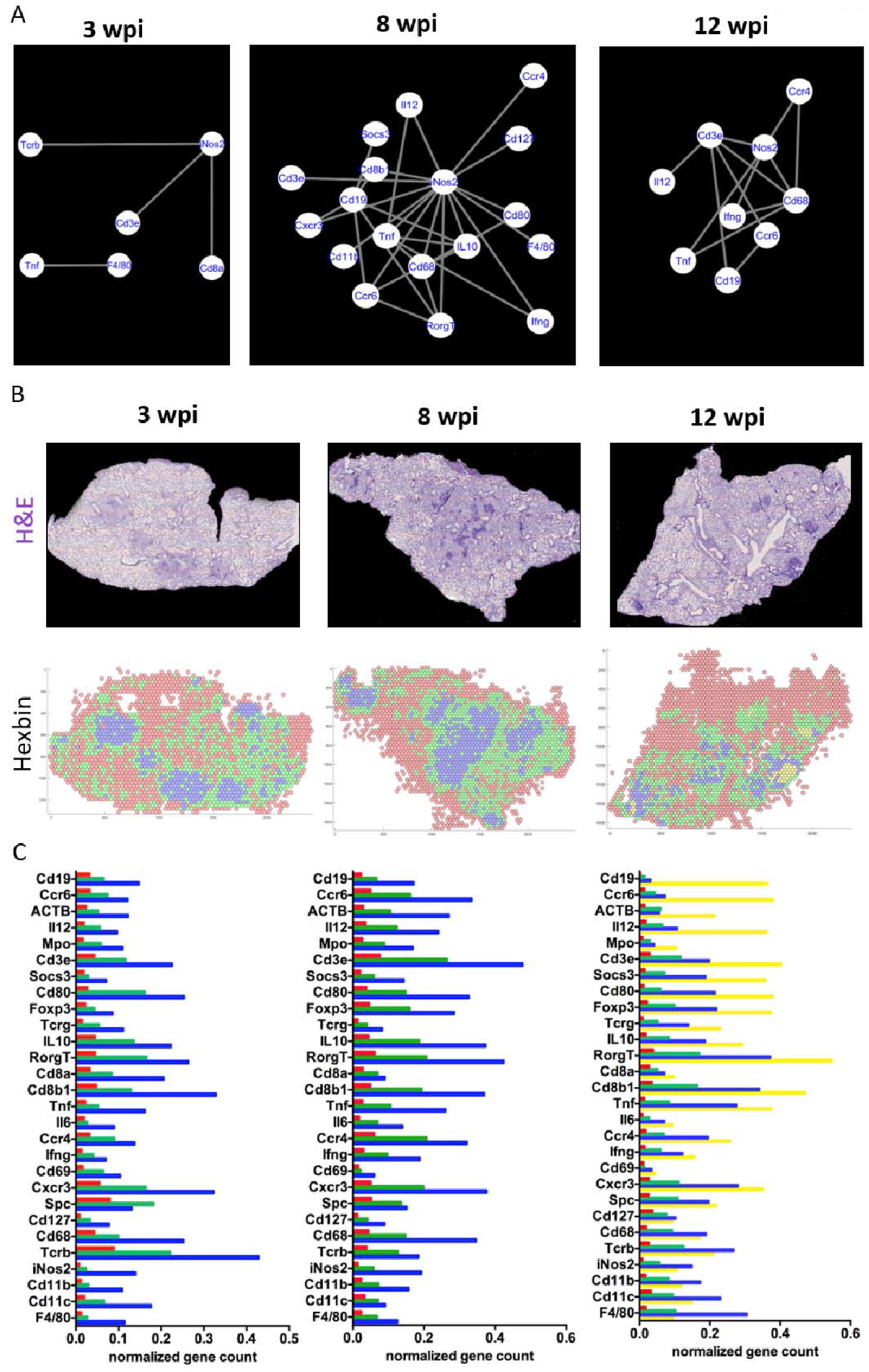
Maturation of the granuloma. A. The spatial co-expression relationships between the *in situ* sequencing data was converted into network-based visualization, where each unique transcript is a node in the network and edges represent the interactions using InsituNet. To identify spatially co-expressed transcripts, InsituNet analyzed the co-occurrence of transcript detections within 30 pixels (10 μm) Euclidean distance for each pairwise combination of transcripts (see Table S1 as example). Representative examples of one lesion per time point as defined in S1 were selected. The significantly co-expressed sequences that are common in lesions from at least two consecutive sections are here depicted. B. The tissue section plane was uniformly tiled into 200 pxls (70 μm) radius hexagons and the density of the multiple sequences in each hexagon was aggregated by binning and displayed into a 2D hexbin map. The densities of the sequences were organized by clustering the hexagons into 3 (3 and 8 wpi) or 4 (12 wpi) different expression patterns. The H&E staining of one representation lung section and their hexbin maps at 3, 8 and 12 wpi are shown. C. The mean centroid normalized transcript counts in each hexagon was compared for the clusters. Note that the red clusters corresponding to unaffected areas contained less counts for of all specific sequences. Note also at 12 wpi that while some sequences were dominant in single clusters (i.e. *cd19* mRNA in the yellow cluster) other sequences were more evenly distributed, or predominated in the blue clusters.

We observed that granulomas, showed co-expression of *inos* and different T cell-related transcripts (*cd3e, tcrb, cd8*) at 3 wpi (Figure 2A). *Inos* sequences were also the principal interacting node in granulomas at 8 and 12 wpi, when the number of co-localizing transcripts in the granulomas increased (Figure 2A). No transcript co-expression was detected in unaffected regions of the sections.

### Distinct localization of transcripts within the granuloma

An unsupervised clustering of mRNA densities across the pulmonary tissue was then performed. The tissues were divided into hexagons (hexbin) with a 70 μm long radius (Figure 2B). The hexbins were separated into the minimal number of clusters showing different sequence frequencies (3 clusters for 3 and 8 wpi, 4 clusters 12 wpi). The hexbin clusters identified areas that corresponded either to the granulomas or to unaffected sites (as defined by H&E staining, Figure S1) at all times of infection (Figure 2B). At 3 wpi clusters differed by the concentration but not on the relative densities of transcripts (Figure 2B, C). One cluster (blue) located in the granuloma, other (red) in unaffected areas, and the third (green) around the granuloma in areas with mild inflammation or unaffected (Figure 2B and S1). In contrast, at 12 wpi the clusters contained different relative frequencies of transcripts (Figure 2B, C). Three of the 4 hexbin clusters located in the granuloma area. One of these (green) also localized in areas surrounding the granuloma (Figure 2B). Another cluster (yellow) overlapped with lymphoid rich areas within the granuloma and showed an over-representation of *cd19* mRNA, expressed by B cells, and several other transcripts (*ccr6, il12, mpo, cd3*) (Figure 2C). However, myeloid markers (*cd68, inos, cd11b, cd11c*) were mainly expressed in the blue cluster. Thus, distinct areas within the granulomas containing different densities of transcripts were revealed by an unbiased analysis of the images. This indicated an increased compartmentalization of granulomas at late time points after infection.

*Cd19* mRNA showed a sparse and indiscriminate expression at 3 wpi, but a strong focal distribution in lymphoid-rich areas that resembles inducible B cell follicles^13,14^ within the C57BL/6 granuloma at later time points (Figure 3A). Immunohistochemical labelling of B cells with anti-B220 in these sections resulted in a similar pattern further confirming the specificity of the *in situ* sequencing (Figure 3B).

**Figure 3.**
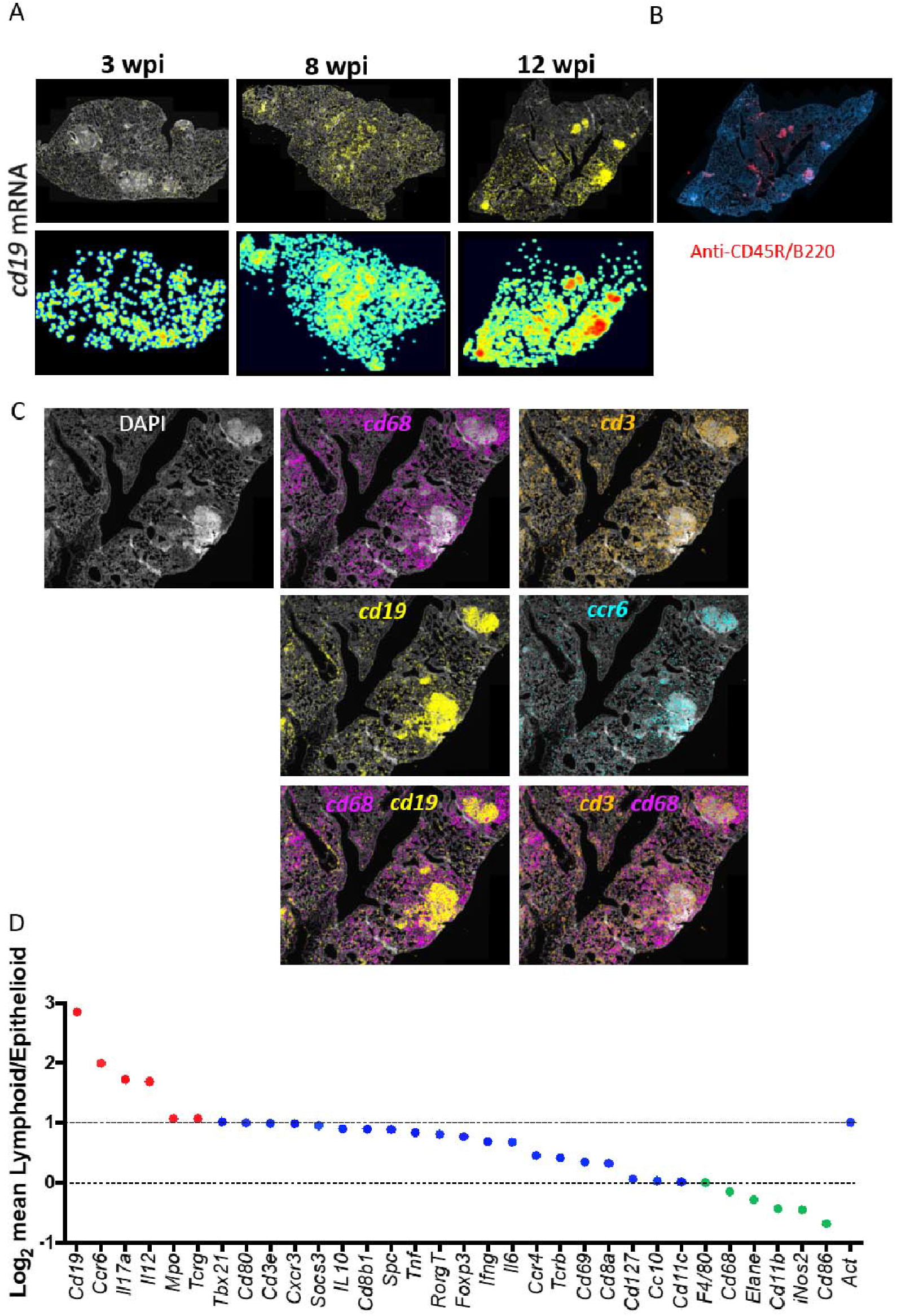
Distinct localization of cd19 mRNA within the lymphoid-rich areas in the granuloma. A. *In situ* detection of *cd19* mRNA transcripts in lungs from *M. tuberculosis*-infected animals. For one representative of 3 consecutive sections per time point the DAPI staining, *cd19* mRNA raw signals and pseudocolor log_2_ density plots are shown. B. Immunohistochemical labelling of CD45R/B220. Note the similar pattern of the labelling as compared to the *in situ* staining for *cd19* mRNA in a consecutive section at 12wpi in A. C. The expression of *cd68, cd3e, cd19* and *ccr6* mRNA in one representative granuloma from C57BL/6-infected mice at 12 wpi is shown. *Cd68* and *cd19* sequences locate in distinct areas of the granuloma, while *cd3e* mRNA locates in both *cd68* and *cd19* mRNA rich areas. D. The density of sequences in the epithelioid or lymphoid areas as defined in FigS1 were quantified in 3 consecutive sections at 12 wpi. The ratios of sequence densities in lymphoid/ epithelioid areas were calculated per section and the mean is depicted. The ratios were color coded accorded if their frequency was higher (red) to *act*b mRNA, or lower than the frequencies in epithelioid cells (green). Transcripts with ratios of lymphoid/ epithelioid >1 and less than the ration of *actb* mRNA are depicted in blue.

Epithelioid areas in 8-12 wpi granulomas contained a high density of *cd68* mRNA and were devoid of *cd19* mRNA (Figure 3C). Instead, *cd3* mRNA co-localized with both *cd68* and with *cd19* mRNA at the epithelioid and lymphoid areas (Figure 3C). *Inos* and *cd68* mRNA localized in the same areas of the lesions, while other transcripts, such as *tnf* and *il12p35/40* mRNA only partially co-localized with the former ones (Figure S3A). *Ccr6 and cd19* mRNA showed a similar localization (Figure 3C).

The density of several transcripts including *ccr6, cd19, il12* and to a lesser degree *cd3, cxcr3* and *cd8b* mRNA was increased in the lymphoid in relation to the epithelioid areas (Figure 3D) after a supervised annotation of the lymphoid and the epithelioid areas of the granulomas based on H&E staining. *Cc10* and myeloid-associated markers were found among the underrepresented transcripts (Figure 3D). In contrast, *cd11b, inos* and *cd68* mRNA were enhanced in the epithelioid as compared to the lymphoid area, while a third group (including *il6, tnf, ifng, cd127* and *ccr4* mRNA) showed similar densities in both areas (Figure 3D).

Thus, in an unsupervised way or after annotation based on histological features, different areas of the granuloma were defined on the frequencies and distribution of transcripts.

Distinct networks of co-expressed transcripts in lymphoid and epithelioid areas of the granulomas were apparent. In lymphoid areas, *cd19* mRNA was a major node interacting with *cd3e* and *ccr6* mRNA at 8 and 12 wpi (Figure 4A, B). *Il12, ifng, tnf* and *cxcr3* mRNA were also found in lymphoid networks at 12 wpi (Figure 4A). *Mpo* (neutrophil myeloperoxidase) and *rorg* transcripts co-localized with *cd19* mRNA as well (Figure 4A). Thus, transcripts from T and B cells are in close proximity in the lymphoid areas.

**Figure 4.**
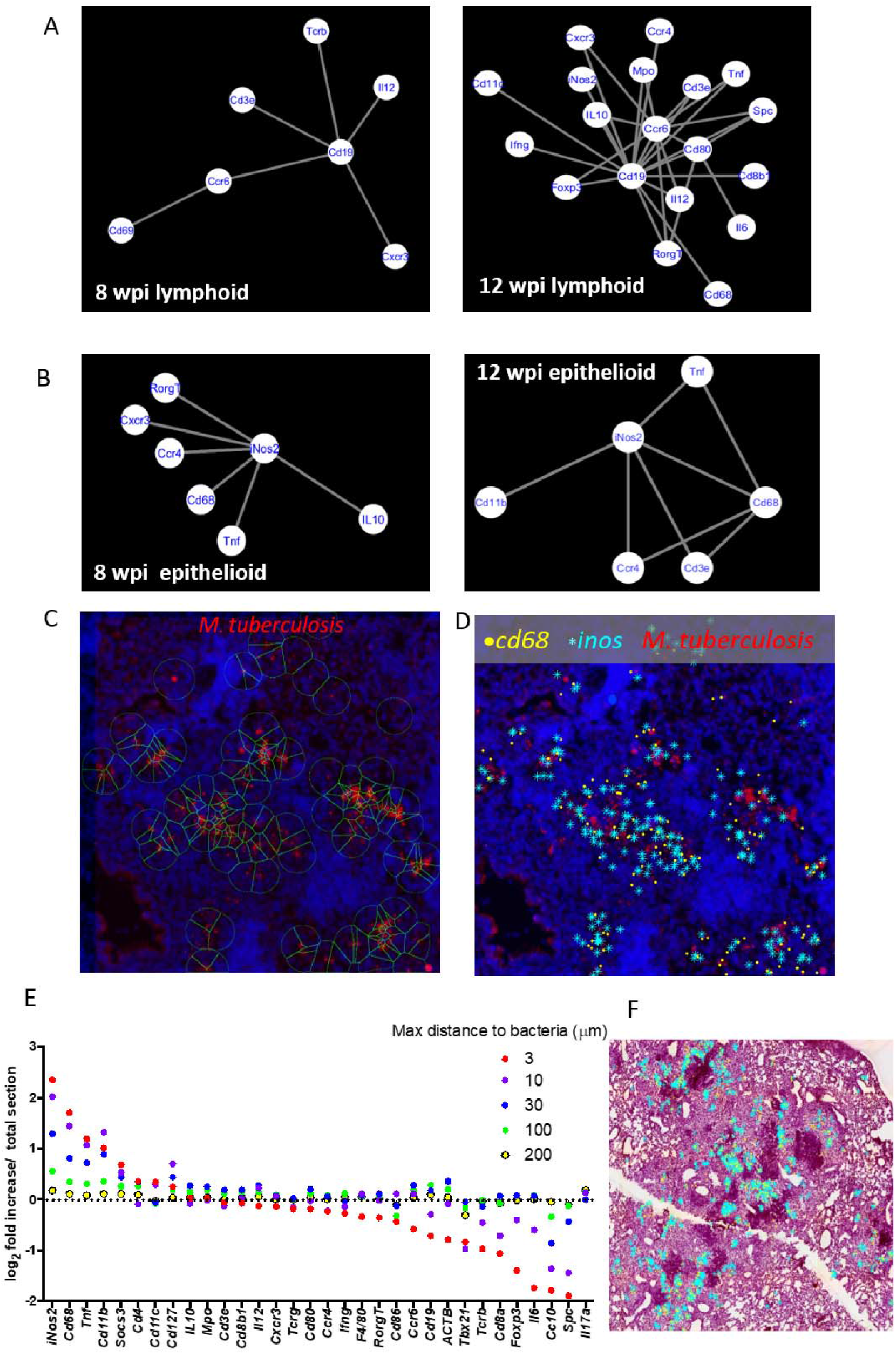
Identification of transcripts co-localizing with M. tuberculosis in tissue sections. A. The networks of co-expressed transcripts in one representative lymphoid area of granulomas at 8 and 12 wpi are shown. B. The networks of co-expressed transcripts in one representative epithelioid area of granulomas at 8 and 12 wpi are shown. C. The auramine-rhodamine-stained *M. tuberculosis* bacteria were aligned with *in situ* transcript signals in tissue sections at 8 wpi. *M. tuberculosis* bacteria were identified by an automated cellprofiler pipeline and a 30 μm radius around those identified is depicted. D. The *inos* and *cd68* transcript signals at < 30μm from identified *M. tuberculosis* bacteria are shown together with *M. tuberculosis* and DAPI labelling for the same selected region as in C. E. The sequences located within a 3, 10, 30, 100 and 200 μm radius from *M. tuberculosis* bacteria were identified. The frequency of each sequence within a given distance was determined in relation to the total transcript count for that distance. The fold increase of this frequency with respect to that observed for the total lung section is depicted. Thus, whether a certain transcript is over- or under represented within the defined distances from *M. tuberculosis* was determined. F. The *cd68* and *inos* transcripts located at < 30 μm are shown aligned with the H&E staining of one representative section (of 3 sections). Note, that most of the extracted sequences are present in the epitheloid region in relative proximity to the lymphoid areas.

On the other hand, *inos*, the main interacting node in the epithelioid area, co-expressed with *cd68* mRNA (Figure 4B). Co-localizing sequences in some epithelioid areas were *ccr4, tnf* as well as other transcripts expressed by T cells (Figure 4B). Density plots of *cd3, cd8b, cxcr3* and *ifng* mRNA showed a similar and wide spread location throughout the granuloma of T cell-related transcripts that only in part overlapped with *ccr4* mRNA (Figure S4A, B).

### Identification of transcripts co-localizing with *M. tuberculosis* in tissue sections

To identify transcripts located in proximity to *M. tuberculosis*, sections were stained with Auramine-Rhodamine T, scanned and aligned to the transcript signals. The transcripts located at less than 3, 10, 30, 300 and 600 μm distance from *M. tuberculosis* bacteria were extracted. *Inos* and *cd68* mRNA at < 10 μm from *M. tuberculosis* are shown (Figure 4 C, D). The frequencies of transcripts localized at expanding distances from the bacteria converged with those in the whole lung, validating the results (Figure S5A, B, Figure 4E). *Inos, cd68, cd11b, tnf* and *socs3* mRNA were enriched at shorter distances to *M. tuberculosis*, suggesting that activated macrophages expressing these transcripts co-localize with bacteria in the granuloma (Figure 4E). Similar results were observed in 3 consecutive slides analyzed as depicted for a selected number of transcripts (Figure S5C). On the other hand, transcripts like *cc10, spc* (expressed by type II alveolar epithelial cells), *il6* and *foxp3* showed an opposite trend: the relative frequency of these transcripts decreased at shorter distances to *M. tuberculosis* bacteria (Figure 4E). Other transcripts including *cd8, cd3e, cxcr3, ccr4* and *il12p40* showed similar frequencies at different distances from *M. tuberculosis*, suggesting a neutral spatial association with the bacteria. Interestingly, *M. tuberculosis* and adjacent transcripts located in the epithelioid areas of the granuloma, in proximity to the lymphoid areas, rather than randomly spread throughout the whole lesion (Figure 4F).

### Heterogeneity of granulomas

The mRNA density in granulomas from the same lung was then compared. Heat maps and principal component analysis showed that 3 out of 4 granulomas showed similar transcript compositions at 3 wpi while a fourth diverged (Figure 5A, C). The three lesions depicted at 12 wpi were heterogeneous (Figure 5B, D). All lesions were different compared to unaffected areas (Figure 5 A-D). The ratio of transcript densities in epithelioid and lymphoid areas of different granulomas within the same lung at 12 wpi also showed substantial variation (Figure 5E).

**Figure 5.**
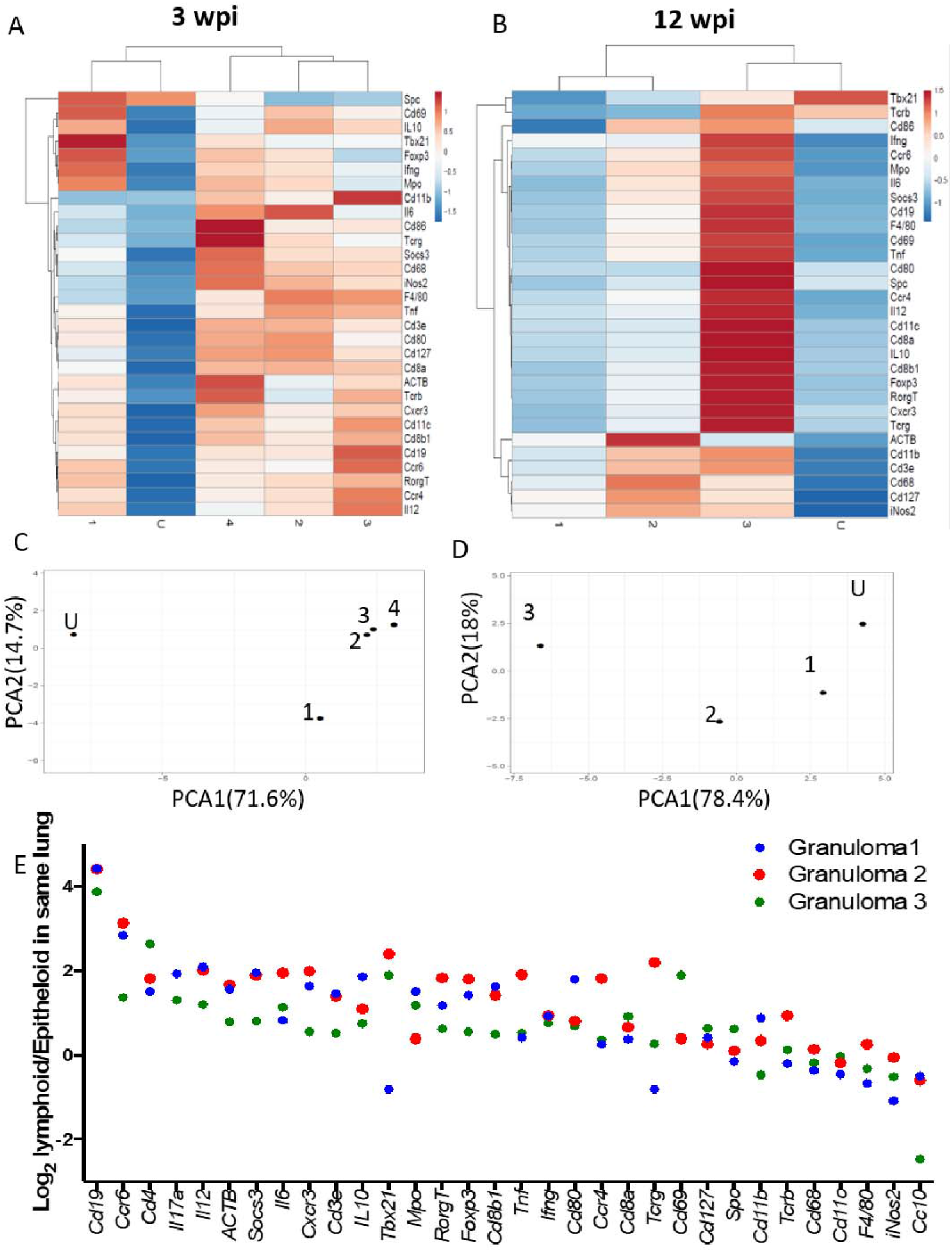
Heterogeneity of granulomas in the same lung. A-D. Heat map analysis depicting the transcript density in different granulomas as defined in S1 at 3 (A) and 12 (B) wpi normalized per area. To simplify the description of the data, results were processed by a multivariate principal component analysis (C, D), showing that while some granulomas cluster together with respect to their transcripts titers, others segregate from these as well as from the unaffected area. One representative result of 3 consecutive sections per time point is shown. E. The density of sequences in lymphoid and epithelioid areas of individual granulomas of one tissue section 12 wpi was extracted. The ratio of sequence density of epithelioid and lymphoid areas per granuloma in the three different granulomas in the same lung was quantified and depicted.

Qualitative differences in networks in lymphoid or epithelioid areas of granulomas from the same lung were also observed (Figure 6A, B). Despite such variation, common networks of transcripts to all epithelioid or lymphoid area determined *inos-cd68* and *cd19* mRNA as the main interacting nodes, respectively (Figure 6C). Those differed substantially from the core networks of necrotic granulomas in the C3HeB/FeJ mice analysed below (Figure 6D).

**Figure 6.**
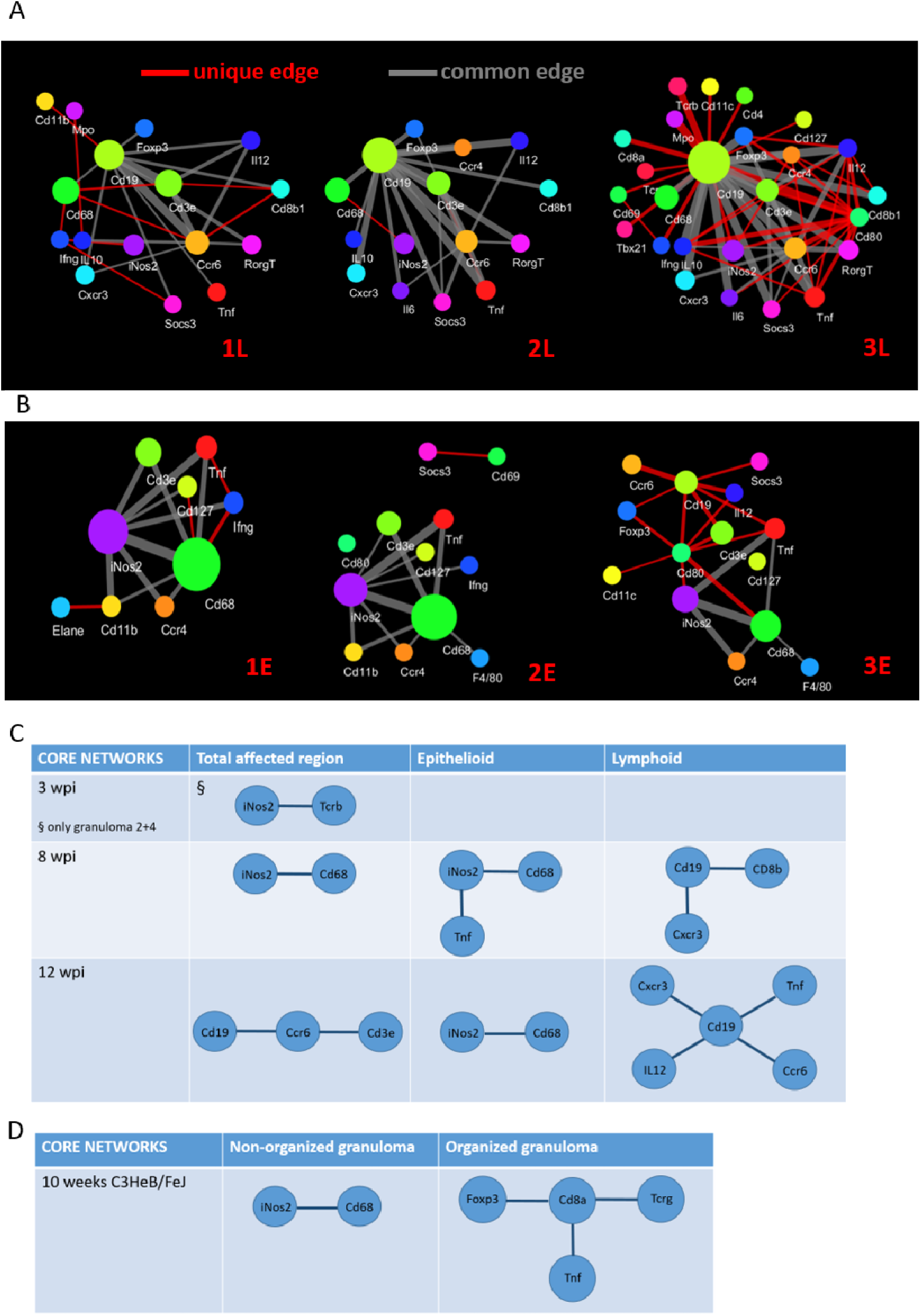
Heterogeneity of transcript networks in areas of the granuloma during infection with M. tuberculosis. A-B. Diversity of interacting networks of transcripts in the lymphoid (A) and epithelioid (B) area of 3 granulomas from one representative lung 12 wpi is depicted. Nodes in the network represent unique transcripts and node size is proportional to the number of transcript detections. Edges represent significant spatial co-expression between transcripts. The more statistically significant the co-expression is, the greater the weight (thickness) of the edge in the network. Connections that occur in at least two of the three networks per region appear in gray, unique connections in red. C-D. Core networks, defined as co-expressed transcripts at <10 μm in consecutive sections found in all annotated regions (Figure S1) for each time point, calculated as described in the method section, are depicted. Diverse core networks were defined for whole granulomas, for epithelioid and lymphoid regions of the granulomas at different time points after *M. tuberculosis* infection in C57BL/6 mice (C). The core networks from non-organized and organized granulomas C3HeB/FeJ mice are also shown (D). No core networks for unaffected regions could be found.

### Comparison of encapsulated and non-encapsulated granulomas

Both encapsulated and non-encapsulated lesions were detected in C3HeB/FeJ *M. tuberculosis*-infected lung sections (Figure 7 A). Highly organized encapsulated granulomas showed a center with caseous and sarcoid necrotic cells surrounded by a fibrotic capsule and a layer of foamy macrophages (Figure S6A). The granuloma contour showed areas with lymphoid and epithelioid cells, as previously described (Figure S6A). *M. tuberculosis* accumulated mainly in the rim of the necrotic granulomas rather than in the center (Figure 7B, S6C) confirming previous data^15^. The center (GC) and the outline (GO) of two large encapsulated granulomas, a non-organized perivascular cellular lesion with high density of lymphocytes and epithelioid cells, with pockets of intracellular bacteria (NOG) (Figure 7A and S6 B, D), a smaller encapsulated granuloma with high bacterial density, areas of necrosis and neutrophilic infiltrates (SG) and a non-affected area were annotated for transcript analysis (Figure S1D). These regions showed different transcript densities. Areas displaying similar morphological features showed related transcript profiles (Figure 7D, E). The center of the encapsulated granulomas (GC) contained a lower *β-actin* mRNA density as compared to other areas (GO, NOG or SG) reflecting the necrotic extension (Figure 7D).

**Figure 7.**
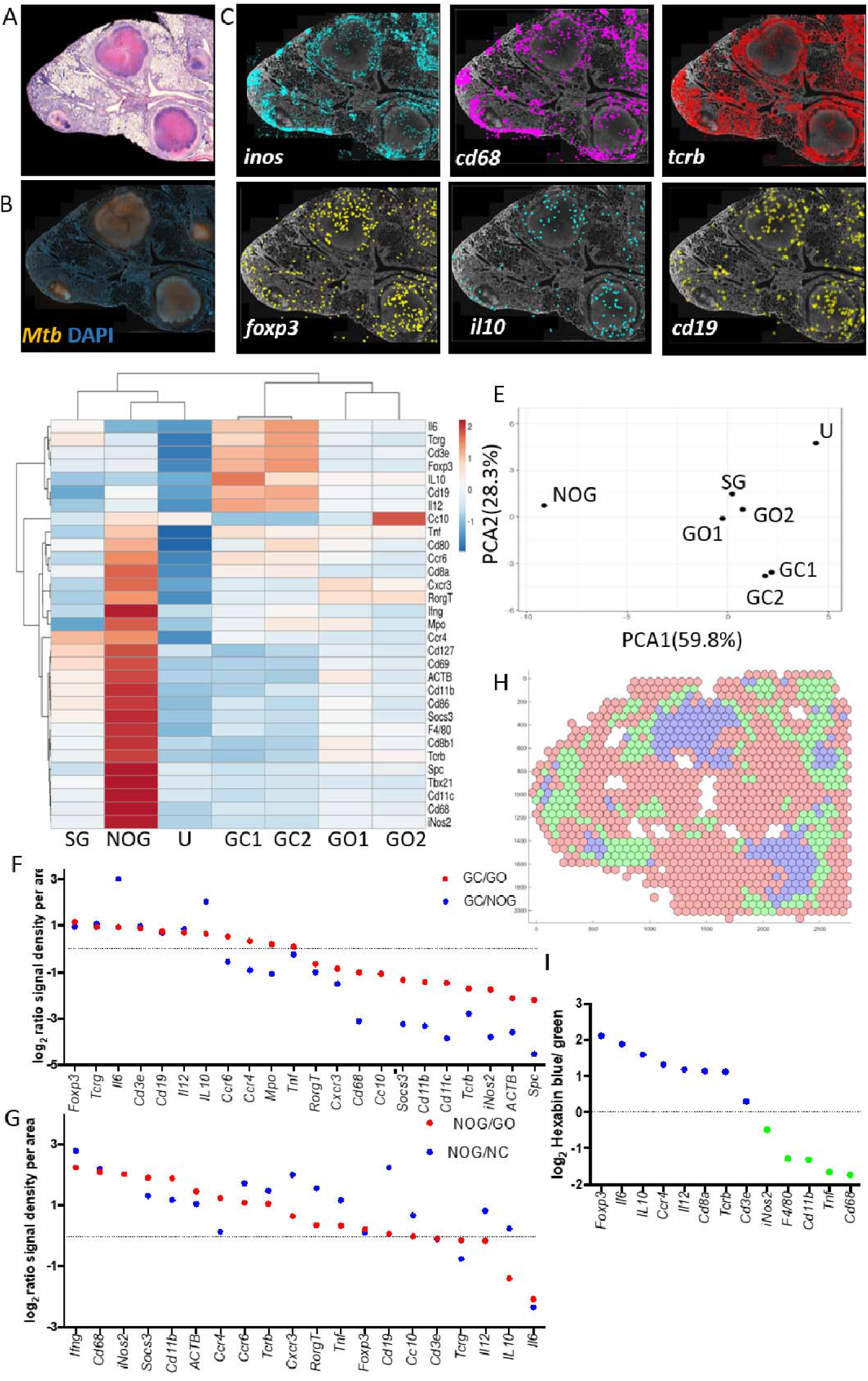
In situ sequencing in encapsulated granulomas. A-B. H&E (A) and auramine-rhodamine-stain of *M. tuberculosis* bacteria aligned with DAPI staining (B) of a C3HeB/FeJ lung section 10 wpi. C. *In situ* signals of *inos, cd68, tcrb, foxp3, il10 and cd19* mRNA transcripts in a lung section from *M. tuberculosis*-infected C3HeB/FeJ, aligned with the DAPI staining, are depicted. D. Heat map analysis depicting the sequence density in annotated areas of a C3HeB/FeJ lung section. The areas correspond to the center (CG) and edge (GO) of encapsulated granulomas, non-encapsulated granulomas with relatively low mycobacterial density (NOG), small-encapsulated granulomas with high bacterial numbers (SG) and an unaffected area (U). E. A multivariate principal component analysis of signals shows proximity between those in annotated areas sharing histopathological features, but the distances between those of different kinds. F. The density of sequences in the different areas of the C3HeB/FeJ lung were extracted. The log2 ratio of sequence density in the center (GC) and edge (GO) of the encapsulated granuloma and in the center vs the non-organized granuloma (NOG) are depicted (D). The correlation between the GC/ CO and GC/ NOG is significant (Pearson p<0.001; r:0.67). G. The log_2_ ratio of sequence density in the non-organized (NOG) vs the surrounding area from the encapsulated granuloma (GO) or the non-controlling granuloma (NC) are depicted. The correlation between the NOG/ GO and NOG/ SG is significant (Pearson p<0.002; r:0.65).

GC contained higher densities of *foxp3* and *il10* mRNA, expressed by regulatory T cells, as compared to the rim or to other regions (Figure 7C-F). GC also showed higher levels of *tcrg* (coding for the γ-chain of the T cell receptor of γδ T cells), *cd3* and *il6* transcripts (Figure 7D, E). Instead, GO showed higher frequencies of *tcrb* (coding for the β-chain of the T cell receptor of αβ T cells), *cxcr3, cd8, cd68* and *inos* mRNA than GC (Figure 7D, F).

Levels of *cd68, inos, cxcr3, tcrb* and *ifng* transcripts were higher in NOG than in the center (GC) or the rim (GO) of the large and the smaller encapsulated granulomas (SG) (Figure 7C, D, G). *Cd19* mRNA clusters were not observed in the lung of *M. tuberculosis*-infected C3HeB/FeJ mice (Figure 7C).

An unbiased hexbin analysis also showed a clear distinction of clusters in their transcript contents that aligned in the necrotic cores (blue) and the surrounding areas of the necrotic and the cellular granuloma (green cluster, Figure 7H). The centroid expression of *foxp3, il6* and *il10* transcripts were highest in the blue cluster in agreement with the analysis of annotated areas (Figure 7I).

A network analysis showed co-expression *foxp3*, *tcrg* and *il6* mRNA in the necrotic granuloma (Figure S6E). Instead, NOG in C3HeB/FeJ mice showed *cd68-inos* interactions resembling interactions described for the epithelioid regions of C57BL/6 granulomas (Figure S6F, 4B).

Altogether, C3HeB/FeJ mice encapsulated granulomas contained a center enriched of transcripts associated with immune suppression while cellular granulomas contained networks associated to protective responses.

## Discussion

Studies using TB granulomas in man came to an end in the 1950s with the introduction of antibiotics and the decline of lung lobectomies^16^. Research interest relocated from morphological histopathology to immunology, molecular microbiology, and genetics studied in isolated cells and animal models. While the understanding of the biology of infection in these areas has increased immensely, that of the granuloma structure has advanced at a slower pace. While the novel methods have recently improved the molecular and cellular dissection of the granuloma, this has largely been done in homogenates or isolated cell suspensions in which the histological context is lost.

Here, the localization of 34 immune transcripts in lungs during infection with *M. tuberculosis* was resolved at cellular resolution and aligned with the tissue topology generating high resolution images of the immunological landscape of granulomas.

### Specificity, reproducibility and performance of *in situ* sequencing

The *in situ* sequencing method displayed a high signal to noise ratio of fluorescent signals and high specificity of analyzed sequences allowing target recognition at the single nucleotide level, as previously reported^9^. One limitation of our study is the small number of independent sections examined. This is due to the extensive image acquisition and data processing. Statistics were performed by comparing frequencies of transcripts in different areas of the lung or in different regions of the granuloma but also by comparison of determinations in consecutive sections, which confirmed the specificity of the signals. The results presented were confirmed in one independent sample for each condition.

### Maturation of the granuloma

Histopathological analyses have determined that the formation of murine and human granulomas is regulated by the orderly recruitment of immune cells^17^. Here, we show that most of the immune transcripts identified located within the granulomas. Whereas the frequency of transcripts expressed by innate immune cells (*tnf, il6, inos, il12* or *cd68*) in the granuloma at different time points after infection was similar, several transcripts expressed by T- or B-cells (*cd3, cd19, ccr6, cxcr3, ifng*) showed an increased relative localization in the granuloma at later time points. Networks of co-expressed transcripts showed an increased complexity of interactions in the granulomas at 8 and 12 compared to 3 wpi. In line with this, the initiation of *M. tuberculosis-specific* T-cell responses that limit bacterial growth has been shown to occur later than immune responses stimulated by other infections^18^.

### Transcript compartmentalization within the granuloma

An unsupervised analysis using clustering estimation of transcript expression patterns in the infected lung represented in hexbin maps showed differences in transcript levels (3 and 8 wpi) early but a distinct relative abundance of specific transcripts in the different clusters was observed only late after infection. At 12 wpi, an overrepresentation of *cd19* mRNA was noted in one of the clusters. *Cd19* mRNA localized in lymphoid cell areas that were detected at 8 and 12, but not 3 weeks after infection. In such lymphoid areas, *cd19* and T cell transcripts were co-expressed together with *ccr6* mRNA in most granulomas, indicating close interactions between T cells and B cells in the lymphoid areas. Notably, CCR6 is expressed on IL-17-expressing cells such as T_H_17 cells and ILC3s and participates in their homing to mucosal microenvironments^19,20^ and RORγt is the key transcription factor required for T_H_17 differentiation and IL-17 expression^21^. CCR6 might also be transiently expressed in germinal center B cells and DCs^22,23^, and has been shown to be essential in the development of lymphoid follicles^24^. Both Th17 cells and B cells are important for the formation of tertiary lymphoid organs^25,26^, including lymphoid aggregates in granulomas during infection with *M. tuberculosis*^27^. T_H_1-associated molecules such as *cxcr3* and *ifng* were also co-expressed in networks of the lymphoid-rich areas.

Ectopic lymphoid tissues or tertiary lymphoid organs are present in tissues during several infections, autoimmune diseases and tumors^28^. Different to ectopic lymphoid tissues in other pathologies^29^ a clear distinction of T cell areas and B follicles is not evident in lungs from *M. tuberculosis* infected C57BL/6 mice. Lymphocytic aggregates in TB granulomas containing proliferating B cells, CXCR5^+^ T cells, FDC networks and high endothelial venules 14 have been described in mice^30^, non-human primates^31^, and humans^13^. Chemokine networks involved in developing lymphoid structures have been associated with protection in experimental infections with influenza and *M. tuberculosis*^14,30^. The presence of these cytokines, and the observed close contact between B cells and T cells, suggests that either specific antigens might be presented to primed T cells, or that naïve T cells may be primed in these areas. In support of this suggestion, mycobacteria tetramer^+^ CD4^+^ and CD8^+^ T cells were found in very high proportions in the lung of mice at different time points of infection while mesenteric lymph nodes were contained very few specific T cells if any (unpublished observation). In several lymphoid areas the expression of *il12p35/40* mRNA was observed. IL-12p40 is a constitutive member of IL-12 and IL-23, cytokines secreted by antigen-presenting cells and required for T_H_1 and T_H_17 differentiation respectively which takes place in the lymph nodes. In line with this, antigen-specific T cells were generated during *M. tuberculosis* infection in lungs from mice devoid of lymph nodes due to the deletion of lymphotoxin-α^32^. Together, the lymphoid-rich areas may constitute the structure where host immune responses to *M. tuberculosis* are orchestrated. This has implications for the design of mucosal vaccines.

T cells also located in the epithelioid areas of the granuloma. In these areas, the main clusters included *cd68, tnf* and *inos* but also *cxcr3* and *ccr4* mRNA expressed by T_H_1 and T_H_2, cells respectively^33^. This suggests both T_H_1 and T_H_2 cells penetrate into the epithelioid compartment. In agreement with the observed compartmentalization, CD68^+^ macrophages in the human tuberculous tissues do not form aggregates with B lymphoid clusters^17^.

Our data indicate the preferential presence of *inos, cd68, tnf* and *cd11b* mRNA sequences at subcellular distances (<3μm) to *M. tuberculosis* and strongly advocate that the lesion contains infected activated macrophages. These transcripts localized in the epithelioid areas of the C57/Bl6 mice granuloma, usually in proximity to the lymphoid regions containing cellular populations that might perpetuate the activation of these inflammatory macrophages. iNOS-derived NO plays a key role in host defense against mycobacteria^34^. A variety of *M. tuberculosis* determinants allow the evasion from macrophage bactericidal mechanisms^35^. Relevant for our data, *M. tuberculosis* can detect oxidative-nitrosative induced changes in the host environment^36^ and respond by producing proteins that limit the toxic effects of these changes, including enzymes that repair NO-damaged bacterial proteins at the proteasome level^37–39^. *M. tuberculosis* also inhibits iNOS recruitment to the phagosome to ensure its intracellular survival^40^. This might explain the mycobacterial localization in TNF- and iNOS-expressing macrophages. Transcripts expressed by airway or bronchiolar epithelial cells were negatively associated with *M. tuberculosis*, despite the interaction of the bacteria with airway and alveolar epithelial cells has been shown^41–43^.

### Heterogeneity of granulomas

Structural, microbiological and immunological heterogeneity of TB granulomas in the same lung has been described in animals and humans^44–46^. Both, heat maps and the coexpression network revealed significant variances in transcript localization among different lesions in the same individual. We also show a significant variance between different lymphoid and epithelioid areas of different granulomas, indicating that the differences are not only due to a diverse ratio between both areas. Thus, each granuloma probably represents a limited microenvironment that might be influenced independently from the others in the same lung by the quality of the local immune response and the level of inflammation. However, common core-networks that are unique for each region, time point and mouse strain were here defined.

### In situ sequencing of encapsulated granulomas

C3HeB/FeJ mice developed encapsulated and non-encapsulated granulomas after *M. tuberculosis* infection also displaying distinct transcript patterns. The encapsulated granuloma contained a hypoxic necrotic central area surrounded by a thick fibrotic capsule separating the lesion from other lung areas^15,47^. The increased levels of *foxp3* and *il10* mRNA found in the encapsulated granuloma center support a local suppression of pulmonary inflammatory cells. The presence and suppressive function of *foxp3* expressing regulatory T cells (T_regs_) in granulomatous responses has been previously shown^48–50^, and we suggest also that the localization of these cells coincides with regions of high bacterial density. Similar to our findings, granulomas from children with tuberculous lymphadenitis from showed increased numbers of CD4+FoxP3+ T cells, while CD8+ T cells surrounded the granuloma^51^.

IL-10 is produced at the site of active TB in humans and mice and in both an inhibitory role of protective immunity by IL-10 has been suggested^52–55^. T_regs_, IFN-γ-secreting CD4+ cells, neutrophils and macrophages have all been shown to produce IL-10 during mouse or human TB ^53,56,57^. In association with lower bacterial levels, non-organized granulomas showed a higher frequency of *tcrb, ifng, inos, cd68* and *cd11b* mRNA, that code for proteins of activated macrophages and T_H_1 cells, as compared to the necrotic granuloma.

A combination of laser microdissection and mass spectrometry the abundance of proteins and lipids in different compartments of granulomas from lungs resected from TB HIV and from rabbits has recently been reported, also suggesting variance and common signatures in different compartments of the granuloma^58^. Our findings emphasize the importance of analyses at the tissue-level rather than in blood cell suspensions or in tissue homogenates from tissues for an accurate link to clinical disease and for a selection of proper determinants of disease progression^4^.

The use of *in situ* sequencing in paraformaldehyde-fixed samples eliminates the risk of handling live TB tissues and enables the study of human TB granulomas from either autopsy libraries or surgical biopsies from all stages of disease, including sections from TB patients with comorbidities such as diabetes, obesity or in immunosuppressed individuals to HIV, cancer treatment or ageing. This will allow understanding molecular differences in the buildup of the granuloma in relation to the clinical status of the patients, providing a high-resolution picture of the TB granuloma in different immunological settings.

## Methods

### Mice

C57BL/6 mice were purchased from Janvier labs, C3HeB/FeJ mice were received from Igor Kramnik (BU, Boston, MA). All mice were housed and handled at the Astrid Fagreus Laboratory, Karolinska Institute, Stockholm, under specific pathogen-free conditions and according to directives and guidelines of the Swedish Board of Agriculture, the Swedish Animal Protection Agency, and the Karolinska Institutet.

### Infection and Infectivity Assay

Mice were infected with approximately 250 *M. tuberculosis* bacteria by aerosol using a nose-only exposure unit (In-tox Products). At the indicated time after infection, mice were sacrificed and lungs extracted and fixed in 4% buffered paraformaldehyde for 24 h. Fixed left lungs of mice experimentally inoculated with *M. tuberculosis* were paraffin-embedded. From each lung sample 10 longitudinal 8 μm sections (along the long axis of the lobe) were obtained and stored at −80°C. Slides were paraffin-removed and dehydrated directly before further processing. Sections were stained with hematoxylin-eosin after the *in situ* sequencing procedure. Sections from C57BL/6 mice 8 wpi were also stained with Auramine-Rhodamine T staining mycobacterial lipids following the instructions of the manufacturer (BD).

### Immunohistochemistry

For immunofluorescent staining of mouse B cells, deparaffinized, rehydrated and demasked sections were blocked in 5% BSA for 30 min, washed and incubated with 1:50 anti-mouse with B220 antibody (clone RA3-6B2) in 1% BSA, 0.3% triton-X PBS for 1h at room temperature. After washing in PBS and incubated for 1 hour at RT with secondary donkey-anti-rat rhodamine red 1:100 (both from BD Pharmingen) in antibody buffer. Slides were then washed and mounted in Vectashield with DAPI.

### In situ sequencing technique

The *in situ* sequencing technique was performed as previously described^9^. Briefly, 8 μm tissue sections from paraformaldehyde-fixed and paraffin-embedded lungs were obtained, mounted on microscope slides (Superfrost Plus) and stored at −80°C until further processing. For *in situ* sequencing, mRNA after partial digestion with pepsin at 37°C for 30 min, washed in PBS, then dehydrated by passing them through 70% and 100% ethanol and air-dryed. Gaskets (SecureSeal Hybridization chambers, Grace Bio-Labs) were glued onto the slides such that they form a sealable reaction chamber enclosing the tissue. The mRNA in such sections was in situ reverse transcribed to cDNA using random decamer primers and primers that partially overlap with the recognition sequence of the padlock probes (Table S2). After reverse transcription, an additional crosslinking step was performed (4% paraformaldehyde at room temperature for 45 min), followed by degradation of the mRNA strand and hybridization of padlock probes to the remaining cDNA strand. A ligase in the reaction mix catalyzes circularization of hybridized padlock probes. Multiple Padlock probes were designed for each of the 34 genes of interest such that they detect non-overlapping, transcript-specific sequences (Table S3). Every set of padlock probes for a given transcript carries a unique 4-base barcode, which is used for identification^9^.

*In situ* sequencing substrates are generated in a rolling circle amplification reaction (RCA), which is primed by the cDNA using circularized padlock probes as template. Resulting rolling circle amplification products (RCPs) are subjected to sequencing by ligation. An AlexaFluor750-conjugated sequencing anchor oligo is hybridized immediately adjacent to the barcode sequences of all RCPs (Table S4). Four nonamer libraries are added to the reaction, each containing random nucleotides and one specific base (A, T, G or C) at a fixed position. The libraries are designed such that every base corresponds to a specifc dye. In an enzymatic DNA ligation reaction, the anchor primer is joined to species of the nonamer pool that best match the barcode. After performing the ligation and imaging, the RCPs are reset by stripping off anchor primer and nonamers and a new cycle is initiated. The process is repeated until all four positions of the barcodes have been interrogated. Imaging was performed in an Axio Imager Z2 epifluorescence microscope (Zeiss) by acquiring Z-stacks of overlapping tiles that together cover the tissue section (10 percent overlap). Image stacks were merged to maximum-intensity projections (Zen software).

### Image analysis

A fully automated image analysis pipeline was performed using CellProfiler (v.2.1.1) calling ImageJ plugins for image registration customized to previously described^9^. Briefly, images from all four sequencing cycles were aligned utilizing their general stain for RCPs saving x and y coordinates as well as fluorescence intensities for each RCP in their four base positions to a .csv file and decoded using a Matlab script. For each RCP and hybridization step, the RCP was assigned the base with the highest intensity. A quality score was extracted from each base, defined as the maximum signal (i.e., intensity of assigned letter) divided by the sum of all signals (letters) for that base that ranges from 0.25 to 1. A value close to 0.25 means poor quality (similar signal for all letters), whereas a value close to 1 means that the signal of the assigned letter is strong above a low background. A fixed threshold of 0.45 that gave around 3% unexpected reads was applied.

For signal visualization, Matlab scripts were used to plot selected transcripts on H&E/DAPI-stained images of the analyzed section and to calculate 2log transformations of kernel density estimations for each mRNA species. The number of transcript signals were extracted from the whole tissue and from manually selected regions based on pathological features (Figure S1) for further analysis.

For an unsupervised analysis of our spatial data, we used a Matlab script that generated k-mean clusters for a given number of clusters and size of hexbins. This iterative learning algorithm aims to find groups in the data by clustering them based on similarities in their transcripts expression levels. Transcript counts in every hexbin were normalized by their maximum counts. For each RNA species a cluster centroid (=mean normalized expression level) was computed. The minimum number of clusters that rendered differential results were analysed.

### Heat map and principal component analysis

Extracted transcript reads were normalized to the area and uploaded to ClustVis, a web tool for visualizing clustering of multivariate data using heatmaps and principal component analysis (https://biit.cs.ut.ee/clustvis/)^59^. ClustVis calculates principal components using one of the methods in the pcaMethods R package and plots heatmaps using heatmap R package (version 0.7.7).

### Colocalization analysis

We used two applications in Cytoscape an open source software platform (http://apps.cytoscape.org). InsituNet, a cytoscape app that converts *in situ* sequencing data (above described .csv file with transcript name, x and y coordinates) into interactive network-based visualizations, where each unique transcript is a node in the network and edges represent the spatial co-expression relationships between transcripts, was applied to identify co-expressed transcripts^12^. Co-expressed transcripts were defined in a range of <10 μm and their statistical significance was assessed by label permutation and corrected for multiple testing by the Boniferroni method. Those networks were imported into DynetAnalyzer to further compare networks and extract core-networks^60^.

### Identification of bacteria and surrounding transcripts

After *in situ* sequencing, sections were stained for bacteria with Auramine-Rhodamine T and also hybridized with the fluorescence-labeled anchor primer for the general stain. As for individual base positions multiple focal plane images were acquired and merged to maximum-intensity projections. Bacteria images were aligned to the corresponding four base position images customizing the CellProfiler pipeline from above. In short, bacteria were identified after subtraction of autofluorescence based on global Otsu thresholding, intensities of RCPs were extracted as before with the additional information whether they were located within the assigned distance (3, 10, 30, 300 and 600 μm) to identified bacteria. Custom Matlab scripts (available upon request) were used for decoding and plotting of transcript signals.

## Acknowledgements

We thank the expert help of the staff of the Astrid Fagreus animal house for this study and the comments from Dolores Gavier Widén. We thank also Chika Yokota for expert technical assistance in situ sequencing.

This study was supported by the Swedish Heart and Lung foundation 2015-17/20140641, the Swedish Research Council 2015-02296 and the Swedish Institute for Internationalization of Research (STINT) 4-1796/2014 to MER, and the Karolinska Institutet (M.E.R. and B.C.).

## Author contributions

Investigation, B.C., T. H. and X.Q.; Writing-Original draft, M.E.R.; Review and editing B.C, M.N., M.E.R., T.H. and I.K.; Conceptualization B.C., M.E.R.; Resources M.N. and I. K.; Funding acquisition, M.E.R.

