## Supplementary figures and tables for "Spatial and temporal localization of immune transcripts defines hallmarks and diversity in the tuberculosis granuloma"

*By*

Berit Carow[1,*], Thomas Hauling[2,*], Xiaoyan Qian[2], Igor Kramnik[3], Mats Nilsson[2] and Martin E Rottenberg[1#]

### Supplementary figures

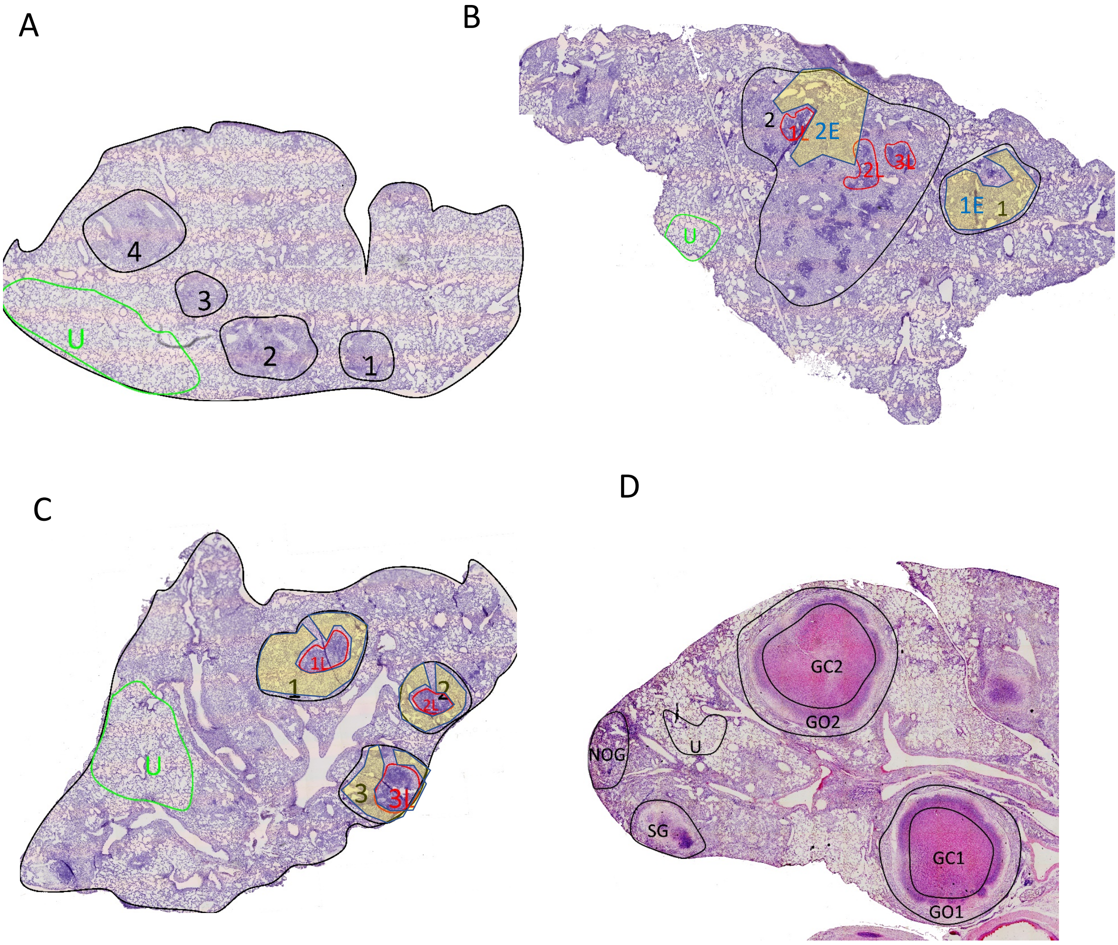


Figure S1

*Histological features of analyzed tissue sections*

H&E stained sections from lung lobes  from C57Bl6 mice at 3 (A), 8 (B), 12 (C) wpi. In D, a section from C3HeB/FeJ mice 10 weeks after infection are shown. Areas that are marked were found and analyzed thoughout the paper for all three consecutive sections. The areas occupied by lesions are marked and numbered in black. Unaffected areas for all lungs are marked in green. In B and C, the lymphoid rich areas are marked in red and epitheloid areas shaded in yellow. In D encapsulated granulomas and non-encapsulated lesions can be observed.

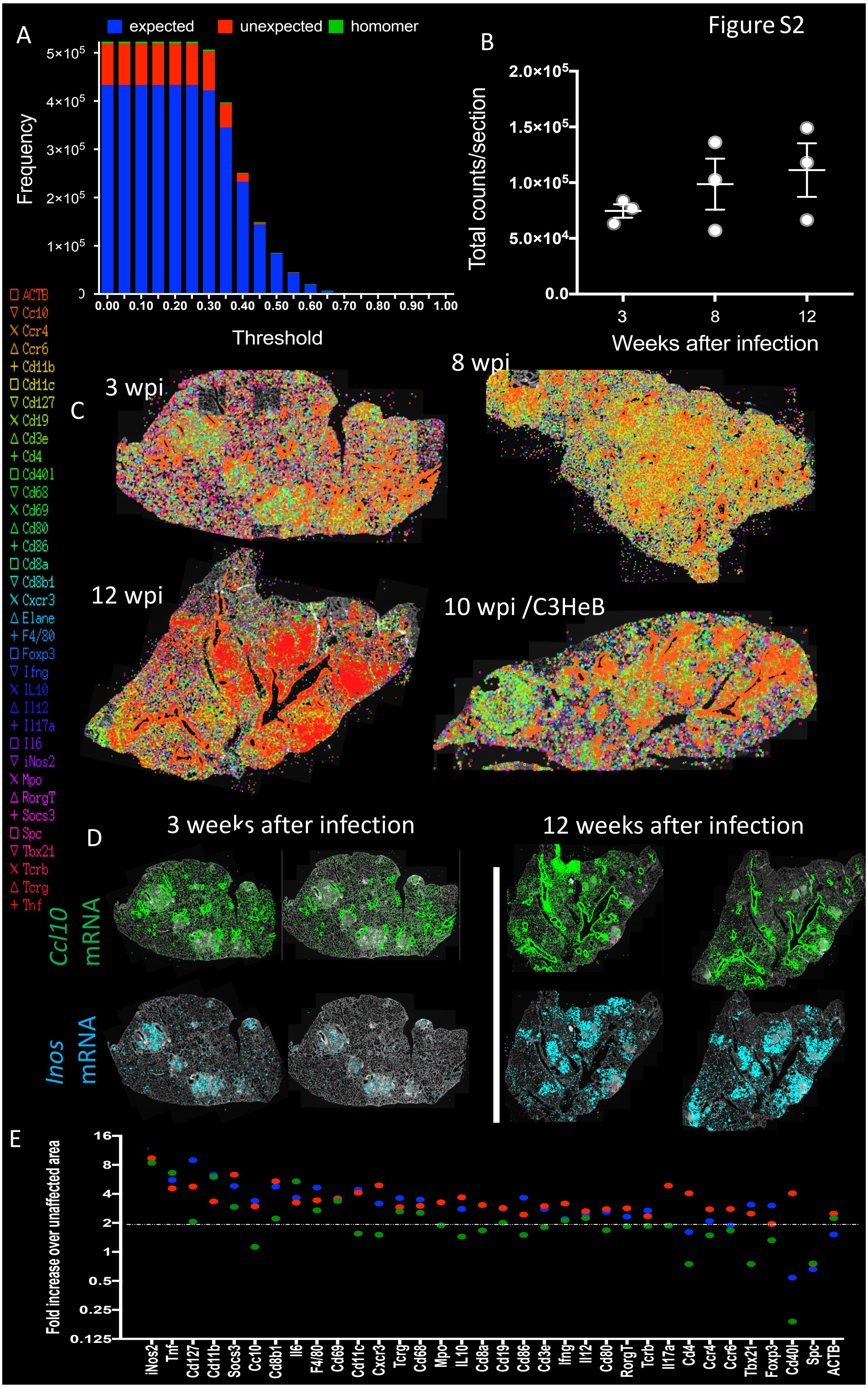
Figure S2
A
expected
unexpected
homomer
Frequency
Threshold
B
Total counts/section
Weeks after infection
C
3 wpi
8 wpi
12 wpi
10 wpi /C3HeB
ACTB
Ccl10
Ccr4
Ccr6
Cd11b
Cd11c
Cd127
Cd19
Cd3e
Cd4
Cd40l
Cd68
Cd69
Cd80
Cd86
Cd8a
Cd8b1
Cxcr3
Elane
F4/80
Foxp3
Ifng
IL10
Il12
Il17a
Il6
iNos2
Mpo
RorgT
Socs3
Spc
Tbx21
Tcrb
Tcrg
Tnf
D
3 weeks after infection
12 weeks after infection
Ccl10 mRNA
Inos mRNA
E
Fold increase over unaffected area

Figure S2

*Specificity and reproducibility of the in situ sequencing*

A. The density of all barcoded sequences in relation to unexpected and homomers (unspecific) reads in each area at different signal intensity thresholds are depicted. The unexpected barcodes showed lower scores than true expected barcodes, and could thus be excluded from further analysis by setting a cut-off at a fixed threshold (0.45).

B. The mean ± SEM of the total number of amplified sequences per section in the different sections analysed are shown.

A. Raw data showing the location of all decoded sequences in paraformaldehyde-fixed lung sections from a *M. tuberculosis* infected mice obtained at the indicated time points after infection. Each dot represent one decoded sequence. The sequences are aligned against a DAPI staining. Note the differential localization of sequences.

B. Raw data showing the location of *cc10* and *inos* decoded transcripts called from two consecutive lung sections from a *M. tuberculosis*-infected mouse obtained at the indicated time points after infection. Each dot represent one decoded sequence. The sequences are aligned against a DAPI staining. Note the differential localization of both sequences and the similarity of consecutive images.

C. The relative fold increase in the density of individual sequences in granuloma vs unaffected lung areas from consecutive lung sections at 3 wpi are shown. A doted line indicates the mean relative density of *actb* mRNA.

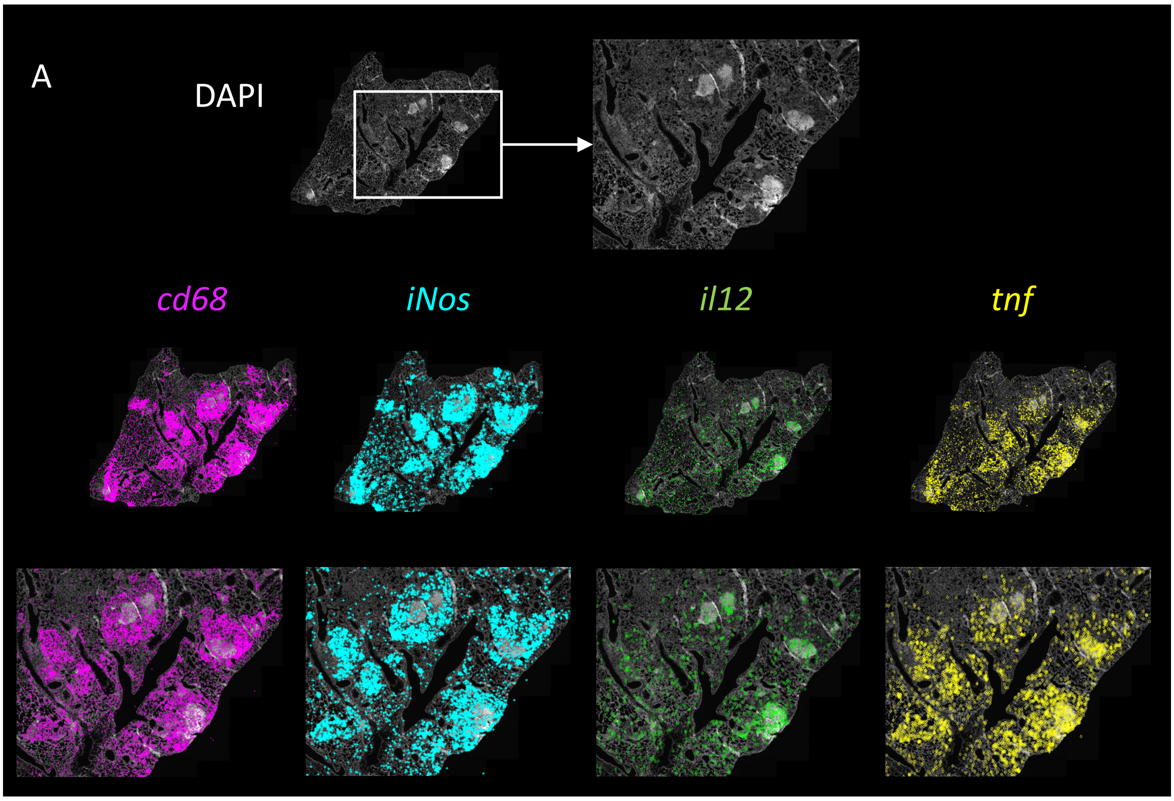


Figure S3

*Spatial distribution of myeloid transcripts in lungs from M. tuberculosis-infected mice*

The raw signals for *tnf, inos, il-12* and *cd68* (A) mRNAs in lungs from *M. tuberculosis* infected mice. Note in A that while *inos* and *cd68* sequences overlap, this is not the case for *cd68* and *il12* transcripts which locate in different areas of the granuloma, better shown in the zoomed area.

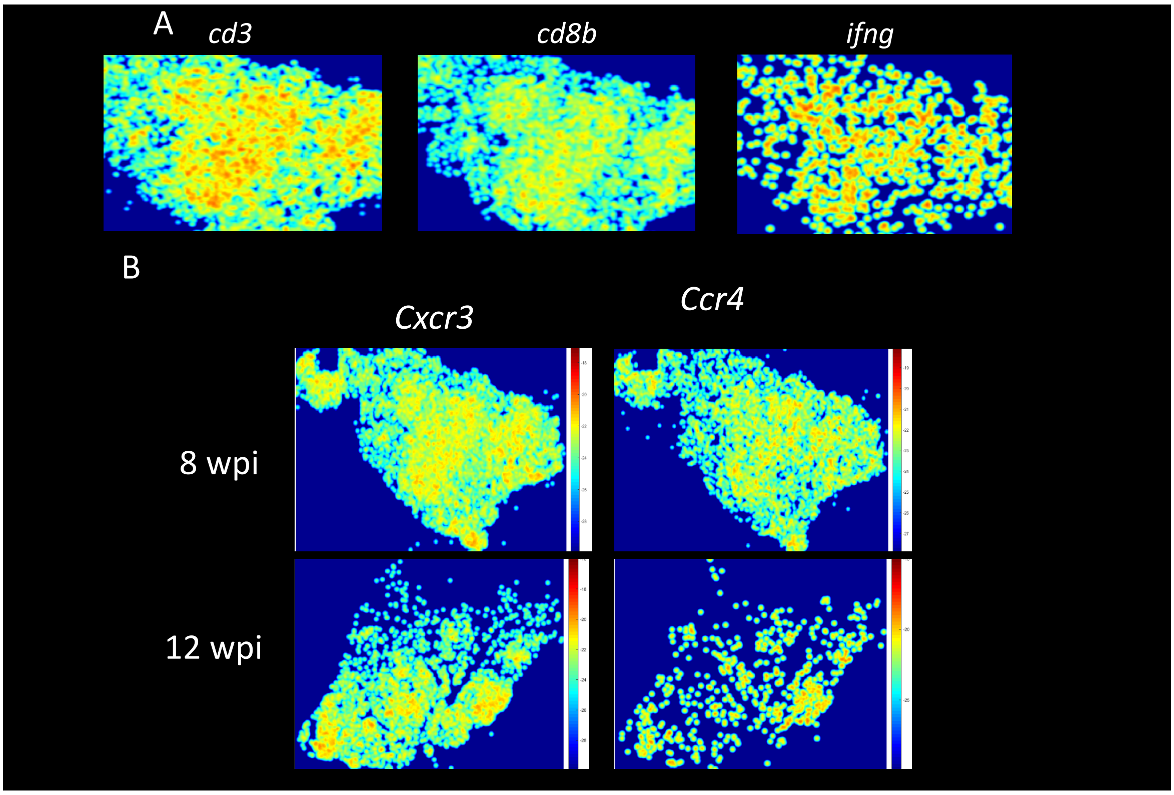


Figure S4

*Spatial distribution of lymphoid transcripts in lungs from M. tuberculosis-infected mice*

A. Pseudocolor log2 density plots of *cd3e, cd8b* and *ifng* mRNA for one representative lung at 8 wpi.

B. Density plot of *cxcr3* and *ccr4* sequences in lungs at 8 and 12 wpi.

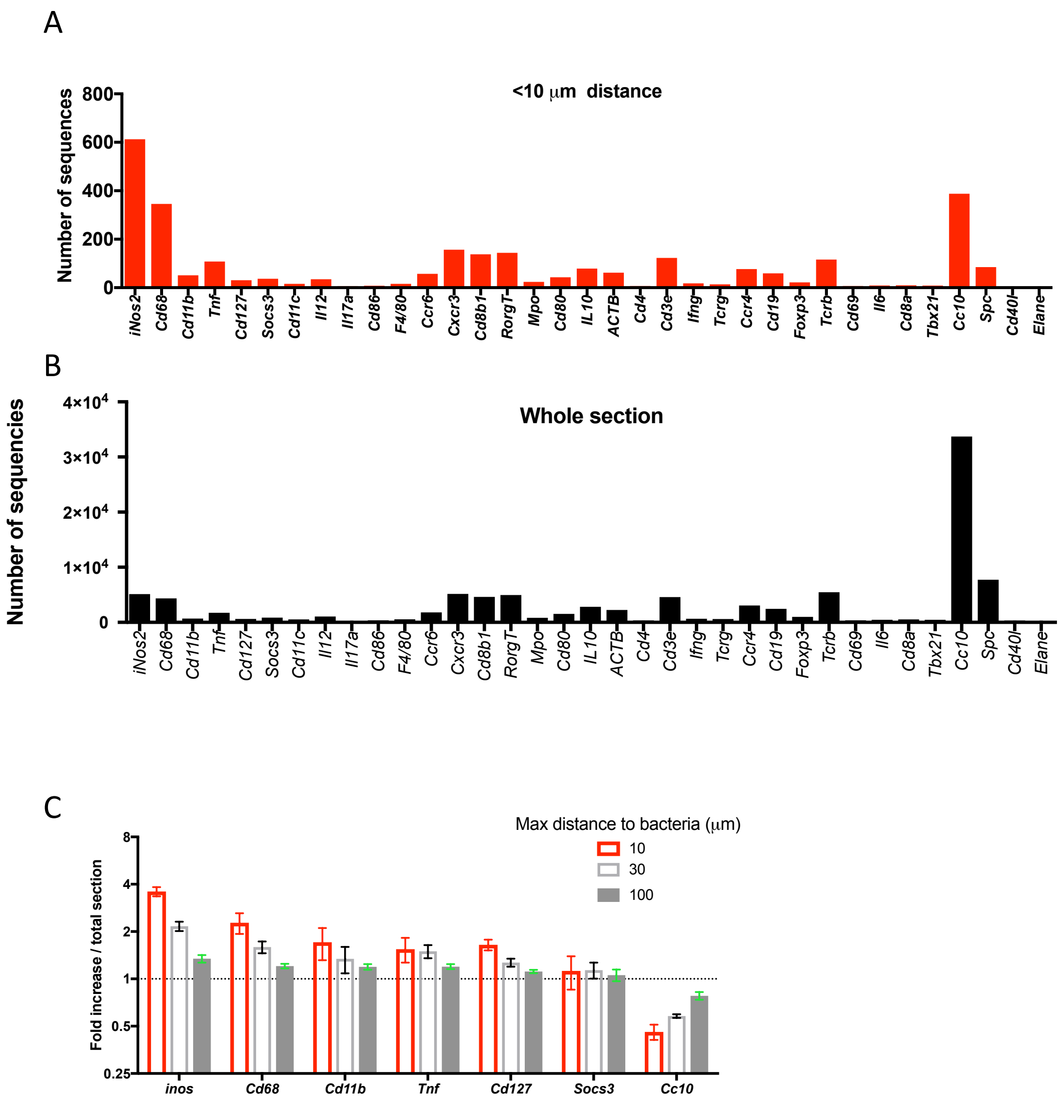


Figure S5

*Identification of sequences expressed at different distances from M. tuberculosis in the lung*

A. The transcript counts for each sequence located at < 10 μm from *M. tuberculosis*.

B. The transcript counts for each sequence in the whole area of the section in (A) are depicted.

C. The frequency of selected transcripts located at <10, 30 or 100 μm from *M. tuberculosis* bacteria in relation to the total sequence count for each distance was determined. The fold increase of such frequencies was compared to the frequencies of each sequence in the whole section. The mean fold increase at each distance vs the whole section of 7 selected transcripts ± SEM of in 3 consecutive sections is depicted.

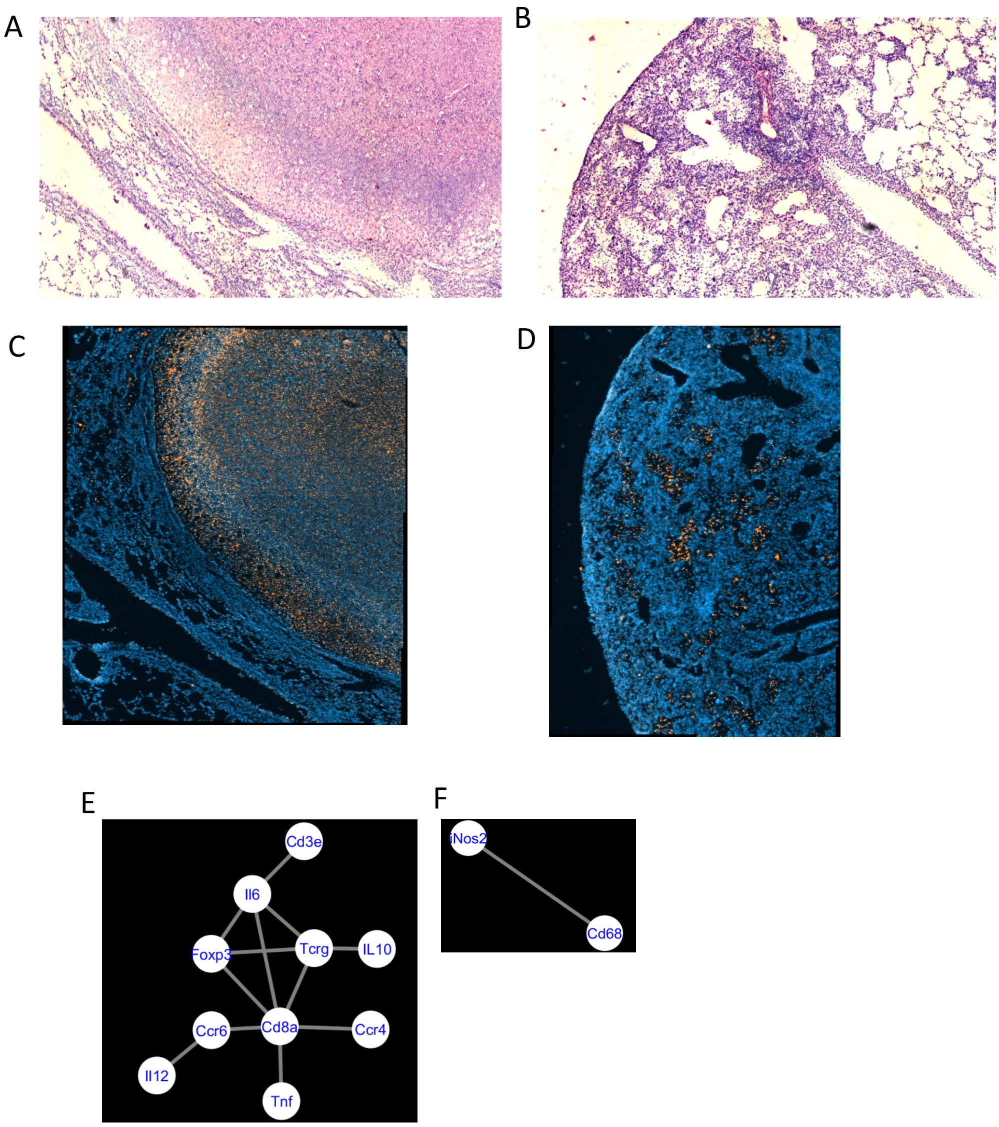


Figure S6

*In situ sequencing of the encapsulated granulomas*

A-B. Micrograph showing the rim area from a necrotic encapsulated granuloma, showing the presence of a capsule, surrounding a layer of epitheloid and foamy cells and delineating compressed lung parenchyma and epitheloid and lymphoid cells (A). Area showing a perivascular lesion with numerous lymphocytes and epitheloid cells in the same H& E stained section from a C3HeB/FeJ lung (B).

C-D. Auramine-rhodamine staining of C3HeB/FeJ encapsulated (C) and non-encapsulated lesions (D). Note bacteria localizes in the rim of the encapsulated lesion.

E-F. The networks of co-expressed transcripts in an encapsulated granuloma (E) and non-encapsulated cellular granuloma (F) are shown.

### Supplementary tables

#### Table S1

*Counts and statistical analysis of paired sequence combinations*

| **Sequences** | **R1L count-section 1** | **p value R1L section 1** | **R1L count section 2** | **p value R1L section 2** |
|---|---|---|---|---|
| *Ccr6-Cd19* | 105 | 0 | 302 | 0 |
| *Ccr6-Cd3e* | 61 | 0 | 126 | 0 |
| *Ccr6-Cd68* | 45 | 8,63E-05 | 99 | 2,36E-07 |
| *Ccr6-Cxcr3* | 16 | 5,46E-05 | 30 | 1,11E-16 |
| *Ccr6-IL10* | 9 | 1,89E-09 | 43 | 0 |
| *Ccr6-Il12* | 14 | 8,88E-16 | 22 | 0 |
| *Ccr6-RorgT* | 20 | 3,65E-08 | 56 | 5,77E-15 |
| *Cd19-Cd3e* | 130 | 0 | 323 | 0 |
| *Cd19-Cd68* | 81 | 3,26E-05 | 229 | 0 |
| *Cd19-Cd8b1* | 25 | 2,35E-12 | 37 | 0 |
| *Cd19-Cxcr3* | 31 | 1,34E-07 | 70 | 0 |
| *Cd19-Il12* | 18 | 5,74E-11 | 67 | 0 |
| *Cd19-RorgT* | 43 | 0 | 106 | 0 |
| *Cd19-Socs3* | 12 | 9,06E-08 | 21 | 0 |
| *Cd19-Tnf* | 27 | 3,84E-05 | 140 | 0 |
| *Cd3e-Cd68* | 100 | 4,33E-05 | 221 | 1,22E-15 |
| *Cd3e-Il12* | 25 | 5,55E-16 | 32 | 0 |
| *Cd68-iNos2* | 146 | 2,33E-09 | 286 | 0 |
| *Ifng-iNos2* | 16 | 3,07E-07 | 22 | 0 |

Example of the counts and p values of statistically significant sequence pairs ($\chi^2$ test) at 10 µm in 1 lymphoid region (R1L see figure S1) of 2 consecutive lung tissue sections at 12 wpi are here shown.

Table S2

*cDNA primers*

| name | sequence |
|---|---|
| mSocs3_prim_2 | CTGGAACTGCCCGGCCGGTC |
| mSocs3_prim_3_AP2 | CTCTCTTGGGGGTACTCCCG |
| mSocs3_prim_4_AP3 | TGGGCTCCAAGATGGCTCAT |
| mMpo_prim_2 | GTGTTGGTTAAACTGAGTG |
| mMpo_prim_3 | AGTTGAGGCCAGTGAAGAAG |
| mMpo_prim_4 | GTGCCAACTCCAGGTTCTTC |
| mIl10_prim_2 | AGTGCTGAGCCAGGCATGAT |
| mIl10_prim_3 | TCAAATTCATTCATGGCCTT |
| mIl10_prim_4 | AAGACCCATGAGTTTCTTCA |
| mIfng_prim_2 | AGGAGGAGAAGCCCAGAACT |
| mIfng_prim_3 | TCCGGCAACAGCTGGTGGAC |
| mIfng_prim_4 | CTCTTGAGACACTGCTTTCT |
| mFoxp3_prim_2 | CATATACCAGGCACAGTGCC |
| mFoxp3_prim_3 | GACCAGGCCGGGAGCACACT |
| mFoxp3_prim_4 | GGCATTGCTTGAGGCTGCGT |
| mCxcr3_prim_2 | CAGGGCGGGGAGTCAGAGAA |
| mCxcr3_prim_3 | GGGGTCCCTGCGGTAGATCT |
| mCxcr3_prim_4 | TGGTCAGAGCGGCCCAGGCG |
| mCd80_prim_2 | AAATGGAACAGAGTGTCTTT |
| mCd80_prim_3 | ATATAAAGTCCGGTTCTTAT |
| mCd80_prim_4 | GACGACGACTGTTATTACTG |
| mCd19_prim_2 | CAGGAAGGGTGTTGACTGGT |
| mCd19_prim_3 | CCCGAGGGAGGCGTCACTTT |
| mCd19_prim_4 | CCCTTGTGGGACACCATGGA |
| mCcr6_prim_2 | AGAACTGTGAAGTTGTTCAC |
| mCcr6_prim_3 | ACTGCCACACAGATGACCTT |
| mCcr6_prim_4 | AGATGTAGCTTTCCGAGTAA |
| mCcr4_prim_2 | CTCCGGGTACCAGCAGGAGA |
| mCcr4_prim_3 | GAAGAGTTGGGTGATGTACT |
| mCcr4_prim_4 | ACTCCTGCCTCTGCCTCCAC |
| mCd11b_prim_2 | GCCAGGTCCATCAAGCCATC |

| | |
|---|---|
| mCd11b_prim_3 | CCAGCATCCTTGTTTTTTAA |
| mCd11b_prim_4 | TGTTTCACATTTCTGCATCA |
| mCd3e_prim_2 | CAGGATGCCCCAGAAAGTGT |
| mCd3e_prim_3 | AGTCTGGGTTGGGAACAGGT |
| mCd3e_prim_4 | AGGAGGTATGGGGTGTGTAA |
| mF4/80_prim_2 | AAGAGGAGCAGCCAAAAGCC |
| mF4/80_prim_3 | GAGCCTGGTACATTGGTGCA |
| mF4/80_prim_4 | ACAGCAGGAAGGTGGCTATG |
| mCd4_prim_2 | CTCGAGACTTTGCAAACAGG |
| mCd4_prim_3 | GTATCTTGAGGGTGAGTGGG |
| mCd4_prim_4 | TCTTGCAAATTCAAAAGAGA |
| mCd8b1_prim_2 | CATCTTGGCGCTTTTGGGGA |
| mCd8b1_prim_3 | CTGGGTCTCTGGGTGGGGGA |
| mCd8b1_prim_4 | TCCAGGGTCCCGGCTAGCTC |
| mCd8a_prim_2 | TGGGGGAGGCGTGTAGGGTC |
| mCd8a_prim_3 | CTAGCTCTGGTGTTACAGTC |
| mCd8a_prim_4 | CAGAAAGAGCCTGGGAATCT |
| mCd11c_prim_2 | CCAGGTACAGCTCATGACTG |
| mCd11c_prim_3 | CTGATTCTCAATATCCTTCA |
| mCd11c_prim_4 | TAGTTGGGTCTTGGGGCTTT |
| mCd127_prim_2 | TATAGCGAAAGCTCTACCCA |
| mCd127_prim_3 | TGAATCTGGCAGTCCAGGAA |
| mCd127_prim_4 | GACAGGTTCATGGCAAGAGA |
| mIl17a_prim_2 | CAGATGAAGCTCTCCCTGGA |
| mIl17a_prim_3 | TGGGGGTTTCTTAGGGGTCA |
| mIl17a_prim_4 | AACATAAAACTAAGTTTGGT |
| miNos2_prim_2 | ATCTCTCCACTGCCCCAGTT |
| miNos2_prim_3 | TCCAGGATGTTGTAGCGCTG |
| miNos2_prim_4 | GTCTAAAGGCTCCGGGCTCT |
| mTcrg_prim_2 | GCAGGAAGTGCTCTGCAGGC |
| mTcrg_prim_3 | ATGACATCGGGAAAGAACTT |
| mTcrg_prim_4 | TTATGGCAGTGAGGATGAAA |
| mRorgT_prim_2 | AGCTCCCGAGATGTCCGGTG |
| mRorgT_prim_3 | ACTTCCTCTGGTAGCTGGTC |
| mRorgT_prim_4 | TGTCAGTCTGTTTTTTATTT |

| | |
|---|---|
| mTcrb_prim_2 | GGGTGCCTGCCGCAAAGTAG |
| mTcrb_prim_3 | TCCAGGGTCCCGGCTAGCTC |
| mTcrb_prim_4 | CTGGGTCTCTGGGTGGGGGA |
| mIl6_prim_3 | TCGTTCTTGGTGGGCTCCAG |
| mIl6_prim_4 | TTGTTCTTCATGTACTCCAG |
| mIl6_prim_5 | ACTTATACATTCCAAGAAAC |
| mIl12_prim_3 | GCAAGGGTGGCCAAAAAGAG |
| mIl12_prim_4 | TTGTCTAGAATGATCTGCTG |
| mIl12_prim_5 | GAGAGAAGCGATGGAGGGGA |
| mCd68_prim_3 | TGCCTTCTCTTGGAAGAGGA |
| mCd68_prim_4 | AGGATTCGGATTTGAATTTG |
| mCd68_prim_5 | AGTGGACTGGGGCAGATGCT |
| mTnf_prim_4 | TTTCTGTTCTCCCTCCTGGC |
| mTnf_prim_5 | AAGAGAACCTGGGAGTAGAC |
| mTnf_prim_6 | CATCTTGTGTTTCTGAGTAG |
| mSCGB1A1_prim | TATCTCTGAAATCCAGTGAG |
| mSftpc_prim | CAGGTCTCTCCCGGAAGAAT |
| Cd69_prim1 | GTGCTTTTGTTTTCTTCCTT |
| Cd69_prim2 | ATCCTTGATGTGATTAGCAG |
| Cd69_prim3 | ATGTCTGATTAGCTTCATTT |
| Cd40lg_prim1 | TCAGATTGTAAGTTCTTAGG |
| Cd40lg_prim2 | CAGTCACGTTGACAAACACA |
| Cd40lg_prim3 | CTTGAGTGTAGACATAATAG |
| mCD86_ap1_prim | CCAGATCTTAAGAGTCTGCA |
| mCD86_ap2_prim | CAAATATACCACTCCCATCC |
| mCD86_ap3_prim | CTGAAGTTGGCGATCACTGA |
| mElane_ap1_prim | CATTATGGCTTCGGATAATG |
| mElane_ap2_prim | TGGGCCACCTGCACGTTGGC |
| mElane_ap3_prim | GCCAGAGTCCGGCTGGATAG |
| mIl12b_ap1_prim | GAGTGTGGCCATTGTGTCCT |
| mIl12b_ap2_prim | GGTTGTCACAAAAGCTAATG |
| mIl12b_ap3_prim | GCACGTGAACCGTCCGGAGT |
| mTbx21_ap1_prim | AGTGATGCAAAACAGAAGAA |
| mTbx21_ap2_prim | CCCTGTTCCTCTGAGGATCC |
| mTbx21_ap3_prim | GTCGGGTCCTGTGCGCCCGG |

| | |
|---|---|
| mCd4_5_prim | CTTGAGGTCTTTGGTGGACT |
| mCd4_6_prim | AGAAGGAGATCCAGGGCTGG |
| mCd4_7_prim | GAACCCTTAGGTTGGACATG |

Table S3

*Padlock probes (all padlock probes are 5' phosphorylated)*

| name | sequence |
|---|---|
| mSocs3_GGAG | TCCAACGTGGCCACCCTCTTCCTCTATGATTACTGACTGCGTCTATTTAG TGGAGCCGGAGCTATCTTCTTTTACTGAGCCGACCTCTCTCC |
| mMpo_GAGG | CGATGACCCCTGCCTCCTCTTCCTCTATGATTACTGACTGCGTCTATTTA GTGGAGCCGAGGCTATCTTCTTTTGCCCTTTGACAGCCTGCA |
| mIl10_CCGG | GCTAACCGACTCCTTAATGCTTCCTCTATGATTACTGACTGCGTCTATTT AGTGGAGCCCCGGCTATCTTCTTTGACCAGCTGGACAACATACT |
| mIfng_CACA | GCTGTTTCTGGCTGTTACTGTTCCTCTATGATTACTGACTGCGTCTATTT AGTGGAGCCCACACTATCTTCTTTCTTTGCAGCTCTTCCTCATG |
| mFoxp3_CAAC | CCTCCCACCACCTTCTGCTTTCCTCTATGATTACTGACTGCGTCTATTTA GTGGAGCCCAACCTATCTTCTTTGACAACCCAGCCATGATCAG |
| mCxcr3_AGAA | GCCCTCTACAGCCTCCTCTTTTCCTCTATGATTACTGACTGCGTCTATTT AGTGGAGCCAGAACTATCTTCTTTTTGACAGAACCTTCCTGCCA |
| mCd80_ACCA | CCGGGGCACATACAGCTGTTTCCTCTATGATTACTGACTGCGTCTATTTA GTGGAGCCACCACTATCTTCTTTCTGGGCCTGGTCCTTTCAGA |
| mCd19_ACAC | GGACTCCTCACCTGTCTCTTTTCCTCTATGATTACTGACTGCGTCTATTT AGTGGAGCCACACCTATCTTCTTTTGTGCTGCCATGCCTCCC |
| mCcr6_AAGA | GTTACTCATGCCACCAACACTTCCTCTATGATTACTGACTGCGTCTATTT AGTGGAGCCAAGACTATCTTCTTTCACCCTACCGTTCTGGGCA |
| mCcr4_AACC | TTTGCTGTTCGTCCTGTCCCTTCCTCTATGATTACTGACTGCGTCTATTTA GTGGAGCCAACCCTATCTTCTTTCTGAACCTGGCCATCTCGGA |
| mCd11b_AATT | AAACCCTAGCCCAAGATTCCTCTATGATTACTGACTGCGTCTATTTAGTG GAGCCAATTCTATCTTCTTTACCTTCAATGACTTCAAGAG |
| mCd3e_ACGT | CACCCTGCTACTCCTTTCCTCTATGATTACTGACTGCGTCTATTTAGTGG AGCCACGTCTATCTTCTTTATCCAGCCCTCCGAG |
| mF4/80_ACTG | TGTCTGCTCAACCGTTCCTCTATGATTACTGACTGCGTCTATTTAGTGGA GCCACTGCTATCTTCTTTCTTTCATCTTCCTCATTCAC |
| mCd4_AGTC | TGCACCGTGACCCTGTCCTCTATGATTACTGACTGCGTCTATTTAGTGGA GCCAGTCCTATCTTCTTTCAGCGACTTCTGGAAC |
| mCd8b1_ATAT | GTTCAAACCAACCATACGTCCTCTATGATTACTGACTGCGTCTATTTAGT GGAGCCATATCTATCTTCTTTCCTTCGTCCCTGCTG |
| mCd8a_ATCG | AAGTGAACTCTACTACTACCTCCTCTATGATTACTGACTGCGTCTATTTA GTGGAGCCATCGCTATCTTCTTTGTGCCAGTCCTTCAGA |
| mCd11c_ATGC | CTGCCACCAACCCTTTCCTCTATGATTACTGACTGCGTCTATTTAGTGGA GCCATGCCTATCTTCTTTGCCTGTCCCTTGCTG |
| mCd127_ATTA | CTTCTCTATTCTTTCTCTCTTCCTCTATGATTACTGACTGCGTCTATTTAG TGGAGCCATTACTATCTTCTTTCACATACAAGCGTGCTT |
| mIl17a_CATG | AGGCAGCCTAAACAGATCCTCTATGATTACTGACTGCGTCTATTTAGTG GAGCCCATGCTATCTTCTTTCCTCGATTGTCCGCC |
| miNos2_GACT | AAGGCCACATCGGATTTCCTCTATGATTACTGACTGCGTCTATTTAGTGG AGCCGACTCTATCTTCTTTCATGACACTCTTCACCAC |
| mTcrg_GCCG | GACCTACCTTTGTCTCCTTCCTCTATGATTACTGACTGCGTCTATTTAGT GGAGCCGCCGCTATCTTCTTTAAATCTCCATAAGGCTGG |

| | |
|---|---|
| mRorgT_GCGC | TCCCTTTCTGCACTCTATTCCTCTATGATTACTGACTGCGTCTATTTAGTGGAGCCGCGCCTATCTTCTTTCCTTCACCCAGCCTT |
| mTcrb_GTAC | GTCTTGTCTGCCACCATCCTCTATGATTACTGACTGCGTCTATTTAGTGGAGCCGTACCTATCTTCTTTCACATCCTATCAACAAGGG |
| mActb_GCAT | TTACACCCTTTCTTTGACAATCCGAGTAGTCTTTGTGCGTCTATTTAGTGGAGCCGCATCTATCTTCTTTGACTGTTACTGAGCTGCGTT |
| mIl6_CGAT_1 | CTCTACGAAGAACTGACAATTCCTCTATGATTACTGACTGCGTCTATTTAGTGGAGCCCGATCTATCTTCTTTGGTATCTGACTTATGTTGTT |
| mIl6_CGAT_2 | GCTCTCCTAACAGATAAGCTTCCTCTATGATTACTGACTGCGTCTATTTAGTGGAGCCCGATCTATCTTCTTTTTCCTACCCCAATTTCCAAT |
| mIl12_CAGT_1 | TGACACCTTTGCTGATTTCTTCCTCTATGATTACTGACTGCGTCTATTTAGTGGAGCCCAGTCTATCTTCTTTCAGTTTTCCTAAGTTCATCA |
| mIl12_CAGT_2 | TCTCCCTCAAGTTCTTTGTTTCCTCTATGATTACTGACTGCGTCTATTTAGTGGAGCCCAGTCTATCTTCTTTGCACTCCCCATTCCTACT |
| mCd68_AGCT_1 | CTGTCTCTCTCATTTCCTTATCCTCTATGATTACTGACTGCGTCTATTTAGTGGAGCCAGCTCTATCTTCTTTGACGGTACCCATCCCCAC |
| mCd68_AGCT_2 | TCCTTCACGATGACACCTTCCTCTATGATTACTGACTGCGTCTATTTAGTGGAGCCAGCTCTATCTTCTTTGCCGTTACTCTCCTGCCA |
| mTnf_CCAA_3 | ATGGCCCAGACCCTCACACTTCCTCTATGATTACTGACTGCGTCTATTTAGTGGAGCCCCAACTATCTTCTTTGCCTCCCTCTCATCAGTTCT |
| mSocs3_GGAG_2_AP1 | GACCGGCCGGGCAGTTCCAGTCTACGATTTTACCAGTGGCTTTTGCGTCTATTTAGTGGAGCCGGAGCTATCTTCTTTGTGACTAAACATTACAAGAA |
| mSocs3_GGAG_3_AP2 | CGGGAGTACCCCCAAGAGAGCTGATTCCTTTGACTCACATTTTTGCGTCTATTTAGTGGAGCCGGAGCTATCTTCTTTCCACCTGCCCAGGCACTCCC |
| mSocs3_GGAG_4_AP3 | ATGAGCCATCTTGGAGCCCATCTACGAGTTTGCAGTCACGTTTTGCGTCTATTTAGTGGAGCCGGAGCTATCTTCTTTGAAGGGAGGCAGATCAACAG |
| mMpo_GAGG_2_AP1 | CACTCAGTTTAACCAACACTCTACGATTTTACCAGTGGCTTTTGCGTCTATTTAGTGGAGCCGAGGCTATCTTCTTTGTTCTTTCAAAGGATTTGGG |
| mMpo_GAGG_3_AP2 | CTTCTTCACTGGCCTCAACTCTGATTCCTTTGACTCACATTTTTGCGTCTATTTAGTGGAGCCGAGGCTATCTTCTTTGAGCCAGCTACCCGGTTCTC |
| mMpo_GAGG_4_AP3 | GAAGAACCTGGAGTTGGCACTCTACGAGTTTGCAGTCACGTTTTGCGTCTATTTAGTGGAGCCGAGGCTATCTTCTTTGGTGAGCTCGGCACGGTGCT |
| mIl10_CCGG_2_AP1 | ATCATGCCTGGCTCAGCACTTCTACGATTTTACCAGTGGCTTTTGCGTCTATTTAGTGGAGCCCCGGCTATCTTCTTTCTTGCAGAAAAGAGAGCTCC |
| mIl10_CCGG_3_AP2 | AAGGCCATGAATGAATTTGACTGATTCCTTTGACTCACATTTTTGCGTCTATTTAGTGGAGCCCCGGCTATCTTCTTTTCCAAGACCAAGGTGTCTAC |
| mIl10_CCGG_4_AP3 | TGAAGAAACTCATGGGTCTTTCTACGAGTTTGCAGTCACGTTTTGCGTCTATTTAGTGGAGCCCCGGCTATCTTCTTTCAGAGCTCCTAAGAGAGTTG |
| mIfng_CACA_2_AP1 | AGTTCTGGGCTTCTCCTCCTTCTACGATTTTACCAGTGGCTTTTGCGTCTATTTAGTGGAGCCCACACTATCTTCTTTACCCTCTGACTTGAGACAGA |
| mIfng_CACA_3_AP2 | GTCCACCAGCTGTTGCCGGACTGATTCCTTTGACTCACATTTTTGCGTCTATTTAGTGGAGCCCACACTATCTTCTTTTCAATGAGCTCATCCGAGTG |
| mIfng_CACA_4_AP3 | AGAAAGCAGTGTCTCAAGAGTCTACGAGTTTGCAGTCACGTTTTGCGTCTATTTAGTGGAGCCCACACTATCTTCTTTAGTAACAGGCTGTCCCTGAA |
| mFoxp3_CAAC_2_AP1 | GGCACTGTGCCTGGTATATGTCTACGATTTTACCAGTGGCTTTTGCGTCTATTTAGTGGAGCCCAACCTATCTTCTTTCTCTGCAGGTTTAGTGCTGT |
| mFoxp3_CAAC_3_AP2 | AGTGTGCTCCCGGCCTGGTCCTGATTCCTTTGACTCACATTTTTGCGTCTATTTAGTGGAGCCCAACCTATCTTCTTTTAGCCACCAGTACTCAGGGC |
| mFoxp3_CAAC_4_AP3 | ACGCAGCCTCAAGCAATGCCTCTACGAGTTTGCAGTCACGTTTTGCGTCTATTTAGTGGAGCCCAACCTATCTTCTTTACACAGGCATAACTGATCAT |

| | |
|---|---|
| mCxcr3_AGAA_2_AP1 | TTCTCTGACTCCCCGCCCTGTCTACGATTTTACCAGTGGCTTTTGCGTCTATTTAGTGGAGCCAGAACTATCTTCTTTATGGGGAAAACGAGAGCGAC |
| mCxcr3_AGAA_3_AP2 | AGATCTACCGCAGGGACCCCCTGATTCCTTTGACTCACATTTTTGCGTCTATTTAGTGGAGCCAGAACTATCTTCTTTGAGCATAGTGCACGCCACCC |
| mCxcr3_AGAA_4_AP3 | CGCCTGGGCCGCTCTGACCATCTACGAGTTTGCAGTCACGTTTTGCGTCTATTTAGTGGAGCCAGAACTATCTTCTTTAAATGTGGATGTTGTTCACG |
| mCd80_ACCA_2_AP1 | AAAGACACTCTGTTCCATTTTCTACGATTTTACCAGTGGCTTTTGCGTCTATTTAGTGGAGCCACCACTATCTTCTTTGGTTGTGAAACTCAACCTTC |
| mCd80_ACCA_3_AP2 | ATAAGAACCGGACTTTATATCTGATTCCTTTGACTCACATTTTTGCGTCTATTTAGTGGAGCCACCACTATCTTCTTTACTAAAAGTGTGGCCCGAGT |
| mCd80_ACCA_4_AP3 | CAGTAATAACAGTCGTCGTCTCTACGAGTTTGCAGTCACGTTTTGCGTCTATTTAGTGGAGCCACCACTATCTTCTTTCTTTGGGGCAGGATTCGGCG |
| mCd19_ACAC_2_AP1 | ACCAGTCAACACCCTTCCTGTCTACGATTTTACCAGTGGCTTTTGCGTCTATTTAGTGGAGCCACACCTATCTTCTTTGCTGGCTTGGTATCGAGGTA |
| mCd19_ACAC_3_AP2 | AAAGTGACGCCTCCCTCGGGCTGATTCCTTTGACTCACATTTTTGCGTCTATTTAGTGGAGCCACACCTATCTTCTTTACCCCGCCAGGAGATTCTTC |
| mCd19_ACAC_4_AP3 | TCCATGGTGTCCCACAAGGGTCTACGAGTTTGCAGTCACGTTTTGCGTCTATTTAGTGGAGCCACACCTATCTTCTTTAATATGTCCAGCCGCTACAT |
| mCcr6_AAGA_2_AP1 | GTGAACAACTTCACAGTTCTTCTACGATTTTACCAGTGGCTTTTGCGTCTATTTAGTGGAGCCAAGACTATCTTCTTTTGCTTCCTGGCCACCGAGGT |
| mCcr6_AAGA_3_AP2 | AAGGTCATCTGTGTGGCAGTCTGATTCCTTTGACTCACATTTTTGCGTCTATTTAGTGGAGCCAAGACTATCTTCTTTCCAGAACACTGACGCACAGT |
| mCcr6_AAGA_4_AP3 | TTACTCGGAAAGCTACATCTTCTACGAGTTTGCAGTCACGTTTTGCGTCTATTTAGTGGAGCCAAGACTATCTTCTTTGGCTTCCTCTGTGCCCGGGT |
| mCcr4_AACC_2_AP1 | TCTCCTGCTGGTACCCGGAGTCTACGATTTTACCAGTGGCTTTTGCGTCTATTTAGTGGAGCCAACCCTATCTTCTTTCTCAACTGTTCTCATTGGCT |
| mCcr4_AACC_3_AP2 | AGTACATCACCCAACTCTTCCTGATTCCTTTGACTCACATTTTTGCGTCTATTTAGTGGAGCCAACCCTATCTTCTTTTCTCGGGGAGAAATTCCGCA |
| mCcr4_AACC_4_AP3 | GTGGAGGCAGAGGCAGGAGTTCTACGAGTTTGCAGTCACGTTTTGCGTCTATTTAGTGGAGCCAACCCTATCTTCTTTGTTATTGGGTATTGGGCATG |
| mCd11b_AATT_2_AP1 | GATGGCTTGATGGACCTGGCTCTACGATTTTACCAGTGGCTTTTGCGTCTATTTAGTGGAGCCAATTCTATCTTCTTTGGGGTAAGGATCTCACAATG |
| mCd11b_AATT_3_AP2 | TTAAAAAACAAGGATGCTGGCTGATTCCTTTGACTCACATTTTTGCGTCTATTTAGTGGAGCCAATTCTATCTTCTTTTTGCATGTCAAGAACAAGTA |
| mCd11b_AATT_4_AP3 | TGATGCAGAAATGTGAAACATCTACGAGTTTGCAGTCACGTTTTGCGTCTATTTAGTGGAGCCAATTCTATCTTCTTTGCGCACCCAGGTCTTTGGAT |
| mCd3e_ACGT_2_AP1 | ACACTTTCTGGGGCATCCTGTCTACGATTTTACCAGTGGCTTTTGCGTCTATTTAGTGGAGCCACGTCTATCTTCTTTTTCTGAGAGGATGCGGTGGA |
| mCd3e_ACGT_3_AP2 | ACCTGTTCCCAACCCAGACTCTGATTCCTTTGACTCACATTTTTGCGTCTATTTAGTGGAGCCACGTCTATCTTCTTTCAAAACAAGGAGCGGCCACC |
| mCd3e_ACGT_4_AP3 | TTACACACCCCATACCTCCTTCTACGAGTTTGCAGTCACGTTTTGCGTCTATTTAGTGGAGCCACGTCTATCTTCTTTCCCGGCCCATCCTCCGACTG |
| mF4/80_ACTG_2_AP1 | GGCTTTTGGCTGCTCCTCTTTCTACGATTTTACCAGTGGCTTTTGCGTCTATTTAGTGGAGCCACTGCTATCTTCTTTACTGCCACAGTACGATGTGG |
| mF4/80_ACTG_3_AP2 | TGCACCAATGTACCAGGCTCCTGATTCCTTTGACTCACATTTTTGCGTCTATTTAGTGGAGCCACTGCTATCTTCTTTAATGTGGACTGAATTCTGTC |
| mF4/80_ACTG_4_AP3 | CATAGCCACCTTCCTGCTGTTCTACGAGTTTGCAGTCACGTTTTGCGTCTATTTAGTGGAGCCACTGCTATCTTCTTTCTGGTATGTCTTGCCTTGGC |
| mCd4_AGTC_2_AP1 | CCTGTTTGCAAAGTCTCGAGTCTACGATTTTACCAGTGGCTTTTGCGTCTATTTAGTGGAGCCAGTCCTATCTTCTTTACTCTCTTCTTCACTAGGTA |

| | |
|---|---|
| mCd4_AGTC_3_AP2 | CCCACTCACCCTCAAGATACCTGATTCCTTTGACTCACATTTTTGCGTCTATTTAGTGGAGCCAGTCCTATCTTCTTTCTCCAGCTGAAGGAAACGCT |
| mCd4_AGTC_4_AP3 | TCTCTTTTGAATTTGCAAGATCTACGAGTTTGCAGTCACGTTTTGCGTCTATTTAGTGGAGCCAGTCCTATCTTCTTTAGCTTCCCATGATGCCTGCT |
| mCd8b1_ATAT_2_AP1 | TCCCCAAAAGCGCCAAGATGTCTACGATTTTACCAGTGGCTTTTGCGTCTATTTAGTGGAGCCATATCTATCTTCTTTGAGCACTGAGGGGAACAGTG |
| mCd8b1_ATAT_3_AP2 | TCCCCCACCCAGAGACCCAGCTGATTCCTTTGACTCACATTTTTGCGTCTATTTAGTGGAGCCATATCTATCTTCTTTGAAGAAGAAGCAATGCCCGT |
| mCd8b1_ATAT_4_AP3 | GAGCTAGCCGGGACCCTGGATCTACGAGTTTGCAGTCACGTTTTGCGTCTATTTAGTGGAGCCATATCTATCTTCTTTCTTTGGCCTTTCATGGAAAA |
| mCd8a_ATCG_2_AP1 | GACCCTACACGCCTCCCCCATCTACGATTTTACCAGTGGCTTTTGCGTCTATTTAGTGGAGCCATCGCTATCTTCTTTGCTAAAGGAGCAGTTTCCCC |
| mCd8a_ATCG_3_AP2 | GACTGTAACACCAGAGCTAGCTGATTCCTTTGACTCACATTTTTGCGTCTATTTAGTGGAGCCATCGCTATCTTCTTTTACAATGGGAGTAATGAGCA |
| mCd8a_ATCG_4_AP3 | AGATTCCCAGGCTCTTTCTGTCTACGAGTTTGCAGTCACGTTTTGCGTCTATTTAGTGGAGCCATCGCTATCTTCTTTTAAAGATTTTTATCTCAGTG |
| mCd11c_ATGC_2_AP1 | CAGTCATGAGCTGTACCTGGTCTACGATTTTACCAGTGGCTTTTGCGTCTATTTAGTGGAGCCATGCCTATCTTCTTTCAGAGCCTGCTTCTGTTCTC |
| mCd11c_ATGC_3_AP2 | TGAAGGATATTGAGAATCAGCTGATTCCTTTGACTCACATTTTTGCGTCTATTTAGTGGAGCCATGCCTATCTTCTTTCGTGGAGAACTTTGATGCTT |
| mCd11c_ATGC_4_AP3 | AAAGCCCCAAGACCCAACTATCTACGAGTTTGCAGTCACGTTTTGCGTCTATTTAGTGGAGCCATGCCTATCTTCTTTTGTCTGCCTTCATATTCATG |
| mCd127_ATTA_2_AP1 | TGGGTAGAGCTTTCGCTATATCTACGATTTTACCAGTGGCTTTTGCGTCTATTTAGTGGAGCCATTACTATCTTCTTTCTCTCTCAGAATGATGGCTC |
| mCd127_ATTA_3_AP2 | TTCCTGGACTGCCAGATTCACTGATTCCTTTGACTCACATTTTTGCGTCTATTTAGTGGAGCCATTACTATCTTCTTTTGAGTTTCAATCCCGAAAGT |
| mCd127_ATTA_4_AP3 | TCTCTTGCCATGAACCTGTCTCTACGAGTTTGCAGTCACGTTTTGCGTCTATTTAGTGGAGCCATTACTATCTTCTTTACACATTCCAACTTCAACCC |
| mIl17a_CATG_2_AP1 | TCCAGGGAGAGCTTCATCTGTCTACGATTTTACCAGTGGCTTTTGCGTCTATTTAGTGGAGCCCATGCTATCTTCTTTTCAGGACGCGCAAACATGAG |
| mIl17a_CATG_3_AP2 | TGACCCCTAAGAAACCCCCACTGATTCCTTTGACTCACATTTTTGCGTCTATTTAGTGGAGCCCATGCTATCTTCTTTCCTAAACAGAGACCCGCGGC |
| mIl17a_CATG_4_AP3 | ACCAAACTTAGTTTTATGTTTCTACGAGTTTGCAGTCACGTTTTGCGTCTATTTAGTGGAGCCCATGCTATCTTCTTTCTCCTCTGAATGGGGTGAAA |
| miNos2_GACT_2_AP1 | AACTGGGGCAGTGGAGAGATTCTACGATTTTACCAGTGGCTTTTGCGTCTATTTAGTGGAGCCGACTCTATCTTCTTTCACAGTATGTGAGGATCAAA |
| miNos2_GACT_3_AP2 | CAGCGCTACAACATCCTGGACTGATTCCTTTGACTCACATTTTTGCGTCTATTTAGTGGAGCCGACTCTATCTTCTTTTTCGAGACTTCTGTGACACA |
| miNos2_GACT_4_AP3 | AGAGCCCGGAGCCTTTAGACTCTACGAGTTTGCAGTCACGTTTTGCGTCTATTTAGTGGAGCCGACTCTATCTTCTTTACAGCAATATAGGCTCATCC |
| mTcrg_GCCG_2_AP1 | GCCTGCAGAGCACTTCCTGCTCTACGATTTTACCAGTGGCTTTTGCGTCTATTTAGTGGAGCCGCCGCTATCTTCTTTGCTGGTGACCTGAAATTCCA |
| mTcrg_GCCG_3_AP2 | AAGTTCTTTCCCGATGTCATCTGATTCCTTTGACTCACATTTTTGCGTCTATTTAGTGGAGCCGCCGCTATCTTCTTTCATACCTTTGTCTCCTTGAA |
| mTcrg_GCCG_4_AP3 | TTTCATCCTCACTGCCATAATCTACGAGTTTGCAGTCACGTTTTGCGTCTATTTAGTGGAGCCGCCGCTATCTTCTTTGGTACAGCAAGTCAGCTGGA |
| mRorgT_GCGC_2_AP1 | CACCGGACATCTCGGGAGCTTCTACGATTTTACCAGTGGCTTTTGCGTCTATTTAGTGGAGCCGCGCCTATCTTCTTTACAGGGCCCCACAGAGACAC |
| mRorgT_GCGC_3_AP2 | GACCAGCTACCAGAGGAAGTCTGATTCCTTTGACTCACATTTTTGCGTCTATTTAGTGGAGCCGCGCCTATCTTCTTTCTCTTTTCACGGGAGGAGGT |

| | |
|---|---|
| mRorgT_GCGC_4_AP3 | AAATAAAAAACAGACTGACATCTACGAGTTTGCAGTCACGTTTTGCGTCTATTTAGTGGAGCCGCGCCTATCTTCTTTAGAGATAGGATGACCAAGTC |
| mTcrb_GTAC_2_AP1 | CTACTTTGCGGCAGGCACCCTCTACGATTTTACCAGTGGCTTTTGCGTCTATTTAGTGGAGCCGTACCTATCTTCTTTAGTCCGGTGAATTCGCCCCT |
| mTcrb_GTAC_3_AP2 | AAGTTCTTTCCCGATGTCATCTGATTCCTTTGACTCACATTTTTGCGTCTATTTAGTGGAGCCGTACCTATCTTCTTTCATACCTTTGTCTCCTTGAA |
| mTcrb_GTAC_4_AP3 | TTTCATCCTCACTGCCATAATCTACGAGTTTGCAGTCACGTTTTGCGTCTATTTAGTGGAGCCGTACCTATCTTCTTTGGTACAGCAAGTCAGCTGGA |
| mIl6_CGAT_3_AP1 | CTGGAGCCCACCAAGAACGATCTACGATTTTACCAGTGGCTTTTGCGTCTATTTAGTGGAGCCCGATCTATCTTCTTTTGTCTGTAGCTCATTCTGCT |
| mIl6_CGAT_4_AP2 | CTGGAGTACATGAAGAACAACTGATTCCTTTGACTCACATTTTTGCGTCTATTTAGTGGAGCCCGATCTATCTTCTTTTTCTGGAGTACCATAGCTAC |
| mIl6_CGAT_5_AP3 | GTTTCTTGGAATGTATAAGTTCTACGAGTTTGCAGTCACGTTTTGCGTCTATTTAGTGGAGCCCGATCTATCTTCTTTTTCTGTTACCTAGCCAGATG |
| mIl12_CAGT_3_AP1 | CTCTTTTTGGCCACCCTTGCTCTACGATTTTACCAGTGGCTTTTGCGTCTATTTAGTGGAGCCCAGTCTATCTTCTTTTGTGTCAATCACGCTACCTC |
| mIl12_CAGT_4_AP2 | CAGCAGATCATTCTAGACAACTGATTCCTTTGACTCACATTTTTGCGTCTATTTAGTGGAGCCCAGTCTATCTTCTTTCACTTCAGAATCACAACCAT |
| mIl12_CAGT_5_AP3 | TCCCCTCCATCGCTTCTCTCTCTACGAGTTTGCAGTCACGTTTTGCGTCTATTTAGTGGAGCCCAGTCTATCTTCTTTGCACAGCTACCTCAGCATGG |
| mCd68_AGCT_3_AP1 | TCCTCTTCCAAGAGAAGGCATCTACGATTTTACCAGTGGCTTTTGCGTCTATTTAGTGGAGCCAGCTCTATCTTCTTTGGGAAGTGAGGCTTTTCATT |
| mCd68_AGCT_4_AP2 | CAAATTCAAATCCGAATCCTCTGATTCCTTTGACTCACATTTTTGCGTCTATTTAGTGGAGCCAGCTCTATCTTCTTTCTTGTGTTCAGCTCCAAGCC |
| mCd68_AGCT_5_AP3 | AGCATCTGCCCCAGTCCACTTCTACGAGTTTGCAGTCACGTTTTGCGTCTATTTAGTGGAGCCAGCTCTATCTTCTTTTCAACCTACCAGCCCCTCTG |
| mTnf_CCAA_4_AP1 | GCCAGGAGGGAGAACAGAAATCTACGATTTTACCAGTGGCTTTTGCGTCTATTTAGTGGAGCCCCAACTATCTTCTTTGCGAGGACAGCAAGGGACTA |
| mTnf_CCAA_5_AP2 | GTCTACTCCCAGGTTCTCTTCTGATTCCTTTGACTCACATTTTTGCGTCTATTTAGTGGAGCCCCAACTATCTTCTTTCAGCCGATGGGTTGTACCTT |
| mTnf_CCAA_6_AP3 | CTACTCAGAAACACAAGATGTCTACGAGTTTGCAGTCACGTTTTGCGTCTATTTAGTGGAGCCCCAACTATCTTCTTTGTGCTCAGAGCTTTCAACAA |
| mCd40L_CGTA_1_AP1 | CCTAAGAACTTACAATCTGATCTACGATTTTACCAGTGGCTTTTGCGTCTATTTAGTGGAGCCCGTACTATCTTCTTTAATCCCAAGGGACCCTGCTC |
| mCd40L_CGTA_2_AP2 | TGTGTTTGTCAACGTGACTGCTGATTCCTTTGACTCACATTTTTGCGTCTATTTAGTGGAGCCCGTACTATCTTCTTTGAATTACAAGCTGGTGCTTC |
| mCd40L_CGTA_3_AP3 | CTATTATGTCTACACTCAAGTCTACGAGTTTGCAGTCACGTTTTGCGTCTATTTAGTGGAGCCCGTACTATCTTCTTTACGGTTAAAAGAGAAGGACT |
| mCd69_CTAG_1_AP1 | AAGGAAGAAAACAAAAGCACTCTACGATTTTACCAGTGGCTTTTGCGTCTATTTAGTGGAGCCCTAGCTATCTTCTTTAAACCTCTGTAGCGTATTTC |
| mCd69_CTAG_2_AP2 | CTGCTAATCACATCAAGGATCTGATTCCTTTGACTCACATTTTTGCGTCTATTTAGTGGAGCCCTAGCTATCTTCTTTCTGAAACTGTCACCAACTGA |
| mCd69_CTAG_3_AP3 | AAATGAAGCTAATCAGACATTCTACGAGTTTGCAGTCACGTTTTGCGTCTATTTAGTGGAGCCCTAGCTATCTTCTTTGAACATTGGATTGGGCTGAA |
| mSCGB1A1_GGCC_AP4 | CTCACTGGATTTCAGAGATATGCTTGTGGTAGCAAATATTTTGCGTCTATTTAGTGGAGCCGGCCCTATCTTCTTTAAGCAAGATTTAAGATTCTGAAG |
| mSftpc_CGCG_AP4 | ATTCTTCCGGGAGAGACCTGTGCTTGTGGTAGCAAATATTTTGCGTCTATTTAGTGGAGCCCGCGCTATCTTCTTTTGATACTGGTTCCGAGTCCG |
| mCD86_CTGA_ap1 | TGCAGACTCTTAAGATCTGGTCTACGATTTTACCAGTGGCTTTTGCGTCTATTTAGTGGAGCCCTGACTATCTTCTTTGTCCATCTTGATTCACCCTG |

| | |
|---|---|
| mCD86_CTGA_ap2 | GGATGGGAGTGGTATATTTGCTGATTCCTTTGACTCACATTTTTGCGTCTATTTAGTGGAGCCCTGACTATCTTCTTTGACAGCTACCTCTTCAGTCA |
| mCD86_CTGA_ap3 | TCAGTGATCGCCAACTTCAGTCTACGAGTTTGCAGTCACGTTTTGCGTCTATTTAGTGGAGCCCTGACTATCTTCTTTAACAGACATTAACAGAACTG |
| mElane_GATC_ap1 | CATTATCCGAAGCCATAATGTCTACGATTTTACCAGTGGCTTTTGCGTCTATTTAGTGGAGCCGATCCTATCTTCTTTTTTGCAGATTGGATCAATTC |
| mElane_GATC_ap2 | GCCAACGTGCAGGTGGCCCACTGATTCCTTTGACTCACATTTTTGCGTCTATTTAGTGGAGCCGATCCTATCTTCTTTATGGCTCCGCTACCATTAAC |
| mElane_GATC_ap3 | CTATCCAGCCGGACTCTGGCTCTACGAGTTTGCAGTCACGTTTTGCGTCTATTTAGTGGAGCCGATCCTATCTTCTTTCCACCATGGCCCTTGGCAGA |
| mIl12b_GCTA_ap1 | AGGACACAATGGCCACACTCTCTACGATTTTACCAGTGGCTTTTGCGTCTATTTAGTGGAGCCGCTACTATCTTCTTTGAAGATACTACGGCTACCTC |
| mIl12b_GCTA_ap2 | CATTAGCTTTTGTGACAACCCTGATTCCTTTGACTCACATTTTTGCGTCTATTTAGTGGAGCCGCTACTATCTTCTTTCAATCAGGGCTGCGTAGGTA |
| mIl12b_GCTA_ap3 | ACTCCGGACGGTTCACGTGCTCTACGAGTTTGCAGTCACGTTTTGCGTCTATTTAGTGGAGCCGCTACTATCTTCTTTGAAGTGTGAAGCACCAAATT |
| mTbx21_GTCA_ap1 | TTCTTCTGTTTTGCATCACTTCTACGATTTTACCAGTGGCTTTTGCGTCTATTTAGTGGAGCCGTCACTATCTTCTTTAGAGTGGTGTCTGGATGTAT |
| mTbx21_GTCA_ap2 | GGATCCTCAGAGGAACAGGGCTGATTCCTTTGACTCACATTTTTGCGTCTATTTAGTGGAGCCGTCACTATCTTCTTTTGCCCATGGACCCGGGCCTG |
| mTbx21_GTCA_ap3 | CCGGGCGCACAGGACCCGACTCTACGAGTTTGCAGTCACGTTTTGCGTCTATTTAGTGGAGCCGTCACTATCTTCTTTATCGTTTCTTCTATCCCGAG |
| mCd4_AGTC_5_AP1 | AGTCCACCAAAGACCTCAAGTCTACGATTTTACCAGTGGCTTTTGCGTCTATTTAGTGGAGCCAGTCCTATCTTCTTTCAAAGAGGTGTCCGTACAAA |
| mCd4_AGTC_6_AP2 | CCAGCCCTGGATCTCCTTCTCTGATTCCTTTGACTCACATTTTTGCGTCTATTTAGTGGAGCCAGTCCTATCTTCTTTGCAGAGAAGGATTCTTTCTT |
| mCd4_AGTC_7_AP3 | CATGTCCAACCTAAGGGTTCTCTACGAGTTTGCAGTCACGTTTTGCGTCTATTTAGTGGAGCCAGTCCTATCTTCTTTAGTGGTTCCAAAGTTCTCTC |

Table S4

*Sequencing-by-ligation oligonucleotides*

| name | sequence | 5' modification | 3' modification |
|---|---|---|---|
| Anchor oligo B2_DO_27_U_5AF750 | /5Alex750N/UGCGUCUAUUUAGU GGAGCC | Alexa Fluor 750 | - |
| Cy5_0A_4N | ANNNCTATC | Phosphorylation | Cy5 |
| Cy3_0G_4N | GNNNCTATC | Phosphorylation | Cy3 |
| TR_0C_4N | CNNNCTATC | Phosphorylation | Texas Red |
| AF488_0T_4N | TNNNCTATC | Phosphorylation | Alexa Fluor 488 |
| Cy5_1A_4N | NANNCTATC | Phosphorylation | Cy5 |
| Cy3_1G_4N | NGNNCTATC | Phosphorylation | Cy3 |
| TR_1C_4N | NCNNCTATC | Phosphorylation | Texas Red |
| AF488_1T_4N | NTNNCTATC | Phosphorylation | Alexa Fluor 488 |
| Cy5_2A_4N | NNANCTATC | Phosphorylation | Cy5 |
| Cy3_2G_4N | NNGNCTATC | Phosphorylation | Cy3 |
| TR_2C_4N | NNCNCTATC | Phosphorylation | Texas Red |
| AF488_2T_4N | NNTNCTATC | Phosphorylation | Alexa Fluor 488 |
| Cy5_3A_4N | NNNACTATC | Phosphorylation | Cy5 |
| Cy3_3G_4N | NNNGCTATC | Phosphorylation | Cy3 |
| TR_3C_4N | NNNCCTATC | Phosphorylation | Texas Red |
| AF488_3T_4N | NNNTCTATC | Phosphorylation | Alexa Fluor 488 |
